# Tau hyperphosphorylation impairs cooperative binding to microtubules and perturbs organelle trafficking in neurons

**DOI:** 10.1101/2025.07.31.667882

**Authors:** Daniel Beaudet, Christopher L. Berger, Adam G. Hendricks

## Abstract

Tau, a neuronal microtubule-associated protein (MAP), organizes the axonal cytoskeleton and regulates intracellular transport. Tau hyperphosphorylation is linked to neurodegeneration in tauopathies including Alzheimer’s disease. Tau binds microtubules cooperatively to form cohesive envelopes, which are thought to control access to the microtubule lattice and regulate the activity of motor proteins and other microtubule-associated proteins. However, how disease-related perturbations affect tau dynamics and its function as a selective barrier to intracellular transport remains unclear. Using tau phospho-variants in vitro and in live neurons, we show that tau hyperphosphorylation disrupts cooperative microtubule binding and dysregulates lysosome transport. Hyperphosphorylated tau does not form envelopes, distributes more uniformly along the axon, and dissociates faster from microtubules. Tau weakly inhibits KIF5C motility, but strongly inhibits KIF1A. Hyperphosphorylation reduces KIF5C inhibition but increases KIF1A inhibition by decreasing processivity and accelerating detachment. Consistent with these effects, hyperphosphorylated tau alters lysosome transport in neurons. While phospho-resistant tau inhibits processive lysosome motility, hyperphosphorylated tau weakens tau-mediated regulation of lysosome transport, mimicking tau knockout neurons that exhibit enhanced processivity. Altogether, these findings show that hyperphosphorylation disrupts tau envelopes and impairs lysosome trafficking, likely contributing to early defects in degradative pathways that drive neurodegeneration.

## Introduction

Tauopathies are a class of neurodegenerative diseases characterized by the misregulation and pathological aggregation of tau, a microtubule-associated protein (MAP) highly expressed in neurons (Binder et al., 1985). Tau regulates the axonal cytoskeleton (Kanai et al., 1992; Chen et al., 1992; Panda et al.,2003; Rosenberg et al., 2008; Chung et al., 2015; Biswas and Kalil, 2018), and acts as a selective barrier on microtubules to direct intracellular transport (Vershinin et al., 2007; Dixit et al., 2008; Vershinin et al., 2008; McVicker et al.,2011; Hoeprich et al., 2014; Hoeprich et al., 2017; Chaudhary et al., 2018; Monroy et al., 2018; Tan et al., 2019; Siahaan et al., 2019; Monroy et al., 2020; Beaudet et al., 2024). Tau’s function and microtubule affinity is modulated by alternative splicing (Kellogg et al., 2018; McVicker et al., 2014) and phosphorylation, both of which play key roles in normal development and pathology (Lindwall and Cole, 1984; Mandelkow et al., 1995; Trinczek et al., 1995; Hasegawa et al., 1998; Niewidok et al., 2016; Kanaan and Grabinski, 2021). In Alzheimer’s disease and other tauopathies, tau becomes abnormally hyperphosphorylated, reducing its microtubule affinity and promoting aggregation (Schneider et al., 1999; Cho and Johnson, 2003). While tau hyperphosphorylation is a hallmark of disease, its impact on the microtubule cytoskeleton and axonal transport, and in turn neuronal proteostasis, remains unclear.

Tau exhibits both diffusive and cooperative interactions with microtubules. Cooperativity leads to cohesive envelope-like structures (Dixit et al., 2008; Hinrichs et al., 2012; McVicker et al., 2014; Siahaan et al., 2019; Tan et al., 2019; Siahaan et al., 2022). Tau envelopes act as selective, reversible barriers along microtubules, regulating the access of motor proteins and other MAPs to the microtubule. Tau cooperativity is well-established in purified, in vitro systems. Envelope formation is sensitive to the intramolecular lattice spacing between α-β tubulin. Tau does not form envelopes on GMPCPP-polymerized microtubules, which are irreversibly expanded, mimicking the GTP-state, but readily forms envelopes on compacted GDP-lattices or reversibly expanded taxol-lattices (Dixit et al., 2008; Monroy et al., 2018; Tan et al., 2019; Siahaan et al., 2019; Siahaan et al., 2022). Discontinuous patterning of tau along microtubules, consistent with envelopes, was also observed in live, non-neuronal U-2 OS cells following taxol treatment (Siahaan et al., 2022) or with elevated pH (Siahaan et al., 2026), and by immunofluorescence staining in fixed hippocampal neurons (Tan et al., 2019). While axonal microtubules are considered to be highly stable (Black et al., 1984; Brady et al.,1984; Sahenk and Brady, 1987; Lim et al., 1989; Okabe and Hirokawa, 1990; Baas et al., 1991; Ahmad et al., 1993; Li and Black, 1996; Song et al., 2013), it was an open question how the rate of tubulin turnover, tubulin posttranslational modifications, and the distribution of lattice conformational states throughout the axon might affect the dynamic localization and cooperativity of tau in neurons.

The extent of tau’s influence on motor proteins depends on several factors including *i)* the tau isoform and posttranslational modifications (PTMs), *ii)* the underlying microtubule lattice, and *iii)* the intrinsic properties of the motors themselves. While some motors, such as kinesin-1 and kinesin-3, are strongly inhibited by tau in an isoform- and lattice-dependent manner, others, like kinesin-2 and dynein, can more effectively navigate tau-decorated regions (Dixit et al., 2008; Vershinin et al., 2007; McVicker et al., 2011; Hoeprich et al., 2014; McVicker et al., 2014; Siahaan et al., 2019; Tan et al., 2019; Siahaan et al., 2022). These studies offer important mechanistic insights into how tau regulates individual motor proteins, but questions remain: how do tau’s regulatory effects scale when multiple motors work in coordinated ensembles to transport organelles? Do additional behaviors of tau, like envelope formation, emerge in vivo that alter transport dynamics? What is the impact of tauopathy-related perturbations on motor protein function?

Using in vitro reconstitution and live-cell imaging in iPSC-derived neurons, we find that disease-associated hyperphosphorylation alters tau dynamics on microtubules and impairs intracellular transport. We used phospho-variants of tau, in which 14 disease-associated residues were mutated to glutamate (E14) to mimic hyperphosphorylation or alanine (AP) to prevent phosphorylation. Our results show that WT and AP tau bind microtubules cooperatively, forming envelopes both in vitro and in live neurons. However, E14 tau interacts more diffusely, has altered distribution along the axon, and fails to form envelopes. Hyperphosphorylation also affects tau-mediated kinesin regulation. WT and AP tau mildly inhibit KIF5C motility but increase detachment within envelopes, whereas E14 tau weaken these effects. In contrast, KIF1A is strongly inhibited by either unphosphorylated or phosphorylated tau. In live tau-knockout neurons, lysosomes move more processively than in control neurons. Rescue assays showed that hyperphosphorylation phenocopies tau knockouts, weakening tau-mediated regulation of lysosome transport, resulting in increased processive motility. Combined, our results demonstrate how disease-related hyperphosphorylation weakens the formation of cooperatively-bound tau envelopes on microtubules and perturbs tau-mediated regulation of degradative organelles, which would be expected to cause defects in cargo maturation and protein homeostasis, contributing to neurodegeneration.

## Results

### Hyperphospho-mimetic tau exhibits diffusive microtubule interactions and reduced tau envelope formation in vitro

We set out to examine how Alzheimer’s disease-associated phosphorylation governs tau’s cooperative binding along microtubules and its function in regulating intracellular transport. Tau contains ∼85 predicted phosphorylation sites (Hanger et al., 2009; Chakraborty et al., 2024). Of which > 45 residues were found to be phosphorylated at higher frequencies in post-mortem brain tissue from Alzheimer’s disease patients (Morishima-Kawashima et al., 1995; Hanger et al.,2007; Hanger et al.,2009; Wesseling et al., 2020). Many disease-associated phosphorylated residues cluster within the proline-rich region of the N-terminal projection domain and the C-terminal region adjacent to the microtubule-binding repeats (Fulga et al., 2007; Steinhilb et al., 2007; Hoover et al., 2010; Wesseling et al., 2020). The microtubule binding repeats, together with the proline rich region and the C-terminus, define the minimal domains required for cooperative binding (Tan et al., 2019). Therefore, we first asked how disease-associated phosphorylation within the proline-rich region and C-terminus alters tau’s cooperative interactions with microtubules.

To directly test the effects of phosphorylation at regions regulating cooperativity, we used in vitro reconstitution assays with mammalian-expressed tau phospho-variants. Mutations either mimic or block phosphorylation at 14 serine and threonine residues in the proline-rich and C-terminal regions (Fulga et al., 2007; Steinhilb et al., 2007; Hoover et al., 2010). We compared WT tau with hyperphospho-mimetic (E14) tau, in which these 14 residues were mutated to glutamate, and phospho-resistant (AP) tau, in which they were mutated to non-polar alanine (Fig 1A). We expressed tau phospho-variants in COS-7 cells because of their widespread use for protein expression and their high transfection efficiency, enabling us to achieve micromolar tau concentrations in lysates. Protein expressed in mammalian cells undergo PTMs and are produced by cellular machinery that more closely resembles neuronal systems than bacterial or insect expression systems. A caveat of this approach is that mammalian expression systems, like insect systems, produce tau with variable phosphorylation patterns (Siahaan et al., 2026; Fan et al., 2025). For example, tau expressed in HEK cells is phosphorylated by endogenous kinases at 58 residues, with <15% occupancy, indicating substantial heterogeneity among tau molecules (Fan et al., 2025). Therefore, expression of these phospho-variant constructs were expected to undergo basal phosphorylation by endogenous kinases, potentially at sites overlapping the 14 mutated residues. Accordingly, the relative phosphorylation levels are predicted to increase from AP tau (lowest) to WT tau (intermediate) to E14 tau (highest).

**Figure 1.**
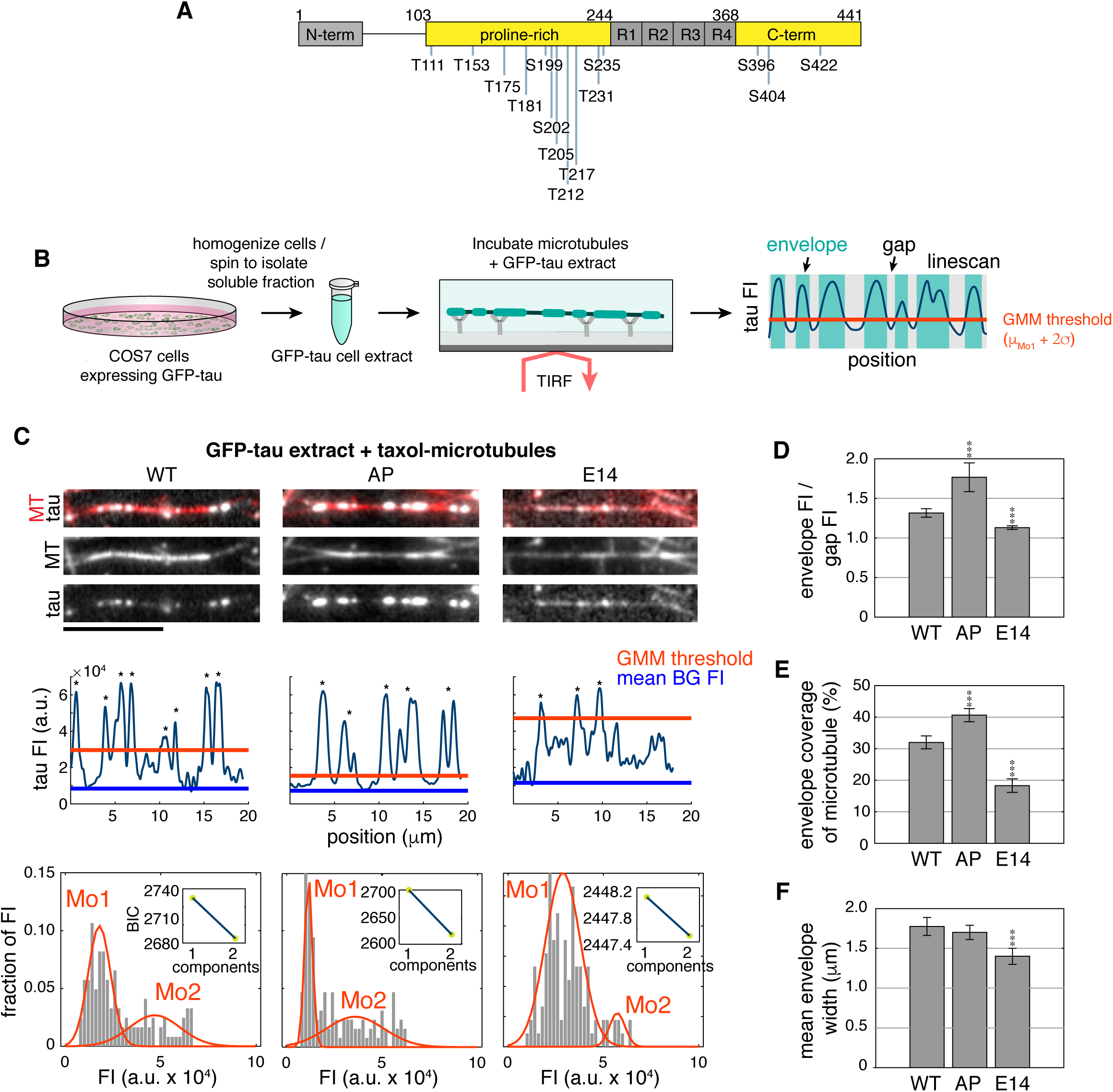
Tau hyperphosphorylation results in more diffuse microtubule interactions and reduces tau envelope formation in vitro. **A)** Structure of 4R0N tau highlighting 14 disease-associated S/T residues mutated to E in pseudo-hyperphosphorylated (E14) tau or to A in phospho-resistant (AP) tau (Hoover et al., 2010). **B)** Schematic illustrating the main steps involved in preparing GFP-tau containing cell extracts and performing reconstitution assays with 500 nM tau (*see* Table 1). Linescans of tau intensity along a microtubule were analyzed using a Gaussian Mixture Model (GMM) to define intensity thresholds distinguishing envelopes from gaps. **C)** Representative images of WT, AP, and E14 GFP-tau (4R0N) on taxol-stabilized microtubules (WT: *n*=183, AP: *n*=170, E14: *n*=170). The indicated *n* values represent total number of samples over 3–4 replicates. Below the images, corresponding tau intensity plots along microtubules are shown, with horizontal lines indicating the GMM threshold (orange) and mean background fluorescence intensity (blue). Asterisks mark peaks identified as envelopes. Histograms show the distributions of tau fluorescence intensity fitted with a GMM. Insets show Bayesian Information Criterion (BIC) analyses used to determine whether a unimodal or multimodal distribution best describes the data. **D–F)** Plots quantifying the effects of hyperphosphorylation on D) tau envelope enrichment intensity, E) percentage of microtubule length covered by envelopes, and F) mean width of envelopes. Error bars indicate 95% CI. Statistical significance was assessed using Student’s t-test (*** p < 0.0001). Scale bars are 10 µm.

We expressed WT, AP, and E14 GFP-tau (4R0N) phospho-variants in COS-7 cells and prepared cell lysates (Fig 1A and B). Lysates were adjusted to a concentration of 500 nM GFP-tau before they were added to flow chambers containing taxol-stabilized microtubules immobilized on glass coverslips (Fig 1B). Our initial observations, consistent with previous studies, showed that tau binds to microtubules in two distinct kinetic phases: cooperative binding that results in the formation of cohesive envelopes and diffusive binding along microtubules (Hinrichs et al., 2012; McVicker et al., 2014; Stern et al., 2017; Tan et al., 2019; Siahaan et al., 2022; Cario et al. 2022; Siahaan et al., 2026; Fan et al., 2025). GFP-tau intensity profiles were generated by manually performing linescans along individual microtubules. The resulting intensity distributions were then analyzed using a non-biased Gaussian Mixture Model (GMM)-based approach to delineate the boundaries between envelopes and gap regions. A threshold based on the first peak of the GMM distribution plus two standard deviations was used to segment tau envelopes, marked by regions of bright pixel intensity that form sharp boundaries with adjacent gap regions containing diffusive tau, characterized by dim pixel intensity (Fig 1B and C). We defined a threshold for each individual microtubule rather than applying a global threshold across microtubules within the same field of view or across replicates. This was necessary due to differences in the fraction of soluble tau between conditions and variations in laser intensity across the TIRF field, both of which introduce variability in tau intensity on microtubules and make a global threshold difficult to apply consistently across replicates. The effects of each phospho-variant were then quantified by measuring envelope coverage along microtubules, envelope width, and tau enrichment in envelopes, which is a measure of the ratio of tau intensity in envelopes relative to gap regions. WT tau formed envelopes with a mean width of 1.77 ± 0.11 μm, covering 32.0 ± 2.0% of microtubules, and an enrichment ratio of 1.32 ± 0.05. AP tau formed more robust envelopes, with a similar mean width of 1.70 ± 0.09 μm, but covering 40.6 ± 2.1% of microtubules, and a higher enrichment of 1.77 ± 0.18. In contrast, E14 tau localized to microtubules but did not show sharp boundaries between envelopes and gaps. E14 tau exhibited the lowest mean envelope width of 1.40 ± 0.10 μm, reduced microtubule coverage of 18.3 ± 2.1%, and a minimal enrichment of 1.13 ± 0.02 (Fig. 1D–F). Thus, phosphorylation strongly reduces the formation, intensity, and frequency of tau envelopes along microtubules.

To examine the effect of hyperphosphorylation on tau-microtubule binding independent of cooperativity, we repeated these experiments using GMPCPP-polymerized microtubules. Previous work showed that tau’s cooperative binding on taxol-stabilized microtubules is facilitated by its ability to compact the spacing between α and β tubulin (Siahaan et al., 2022). Taxol reversibly expands the microtubule lattice, and tau can displace taxol from the microtubule, allowing it to induce lattice compaction. In contrast, GMPCPP microtubules are irreversibly expanded, preventing tau from inducing lattice compaction and thus cooperative binding (Tan et al., 2019; Siahaan et al., 2022). To quantify the effects of phosphorylation on tau’s interaction with microtubules of different lattice states, we measured envelope frequency along GMPCPP- and taxol-stabilized microtubules. We also measured the fraction of bound versus unbound tau by comparing the mean intensity of tau on microtubules to the mean background intensity. As expected, tau did not exhibit significant cooperative binding to GMPCPP microtubules. WT tau displayed an envelope frequency of 0.07 ± 0.02 μm⁻¹ on GMPCPP microtubules versus 0.18 ± 0.01 μm⁻¹ on taxol microtubules, and a lower bound fraction (MT_GMPCPP_: 1.12 ± 0.01 vs to MT_taxol_: 1.32 ± 0.03) (Fig S1B and C). Similarly, AP tau showed a reduced envelope frequency compared to taxol-stabilized microtubules (MT_GMPCPP_: 0.08 ± 0.02 μm⁻¹ vs MT_taxol_: 0.25 ± 0.01 μm⁻¹), along with a reduced bound fraction (MT_GMPCPP_: 1.47 ± 0.05 vs MT_taxol_: 1.85 ± 0.10) (Fig S1B and C). E14 tau envelopes were not detected on GMPCPP microtubules and were significantly reduced on taxol-stabilized microtubules (MT_taxol_: 0.13 ± 0.02 μm⁻¹), as well as showed a reduced bound fraction (MT_GMPCPP_: 1.03 ± 0.01 vs MT_taxol_: 1.11 ± 0.01) (Fig. S1A–C). It is important to note that while a few envelopes were detected for E14 tau using the GMM-approach, they did not form sharp boundaries with gap regions as with WT and AP tau, indicating that E14 tau does not form well-defined envelopes. Together, these results indicate that phosphorylation influences not only cooperativity but also modulates tau’s direct binding to microtubules. Phosphorylation likely also regulates tau-induced lattice compaction (Siahaan et al., 2022), in which misregulation of this process could have downstream effects on microtubule stability, as well as interactions with other MAPs and motor proteins.

### Tau hyperphosphorylation enhances diffusive microtubule interactions and reduces the formation of envelopes in live neurons

Tau binds cooperatively and forms envelopes along microtubules in purified in vitro systems. However, it was unclear whether envelopes form in native neuronal environments and if hyperphosphorylation affects cooperativity in neurons to the same extent as on taxol-stabilized microtubules in vitro. To address this question, we expressed WT, AP, or E14 GFP-tau in CRISPR-edited tau knockout (MAPT-KO) iPSC-derived neurons and imaged them at DIV 7–8, when axons are clearly polarized and synapses formed. This strategy allowed us to directly assess the impact of phosphorylation on tau-microtubule interactions for a single tau isoform, without the added complexity of multiple isoforms and the heterogeneous phosphorylation states as observed with other cells (Siahaan et al., 2026; Fan et al., 2025), both of which significantly influence tau-microtubule interactions. Similar to observations in vitro, tau dispersed either diffusively or formed well-defined envelopes in axons (Fig 2A). Envelopes heterogeneously patterned axons and were often clustered in axonal segments several microns in length between segments of axons with diffuse tau signal (Fig 2A, S2A). To quantify the impact of hyperphosphorylation on envelope formation in neurons, we applied the same GMM-based thresholding method used in our *in vitro* studies (Fig 2A). Quantification showed that WT and AP tau formed envelopes with comparable enrichment intensity (WT: 1.29 ± 0.003 vs AP: 1.32 ± 0.002) and mean width (WT: 2.28 ± 0.22 μm vs AP: 2.27 ± 0.21 μm). In contrast, E14 tau did not form characteristic envelopes with sharp boundaries. Enrichment above thresholds were detected but were much less intense (E14: 1.14 ± 0.002) and the mean width of these areas were reduced compared to WT and AP tau (E14: 1.88 ± 0.24 μm) (Fig. 2B and C). Envelope frequency also varied across conditions. AP tau exhibited a higher frequency of envelopes within discrete regions compared to WT tau (WT: 0.16 ± 0.02 vs AP: 0.22 ± 0.02), as well as a greater percentage of axons contained envelope-positive segments (WT: 35.6 ± 4.5% vs AP: 47.9 ± 4.3%) (Fig 2D and E). In contrast, E14 tau had reduced enrichment frequencies within discrete regions (E14: 0.11 ± 0.03), and these regions were rarely observed throughout axons (E14: 20.3 ± 4.9%) (Fig 2D and E).

**Figure 2.**
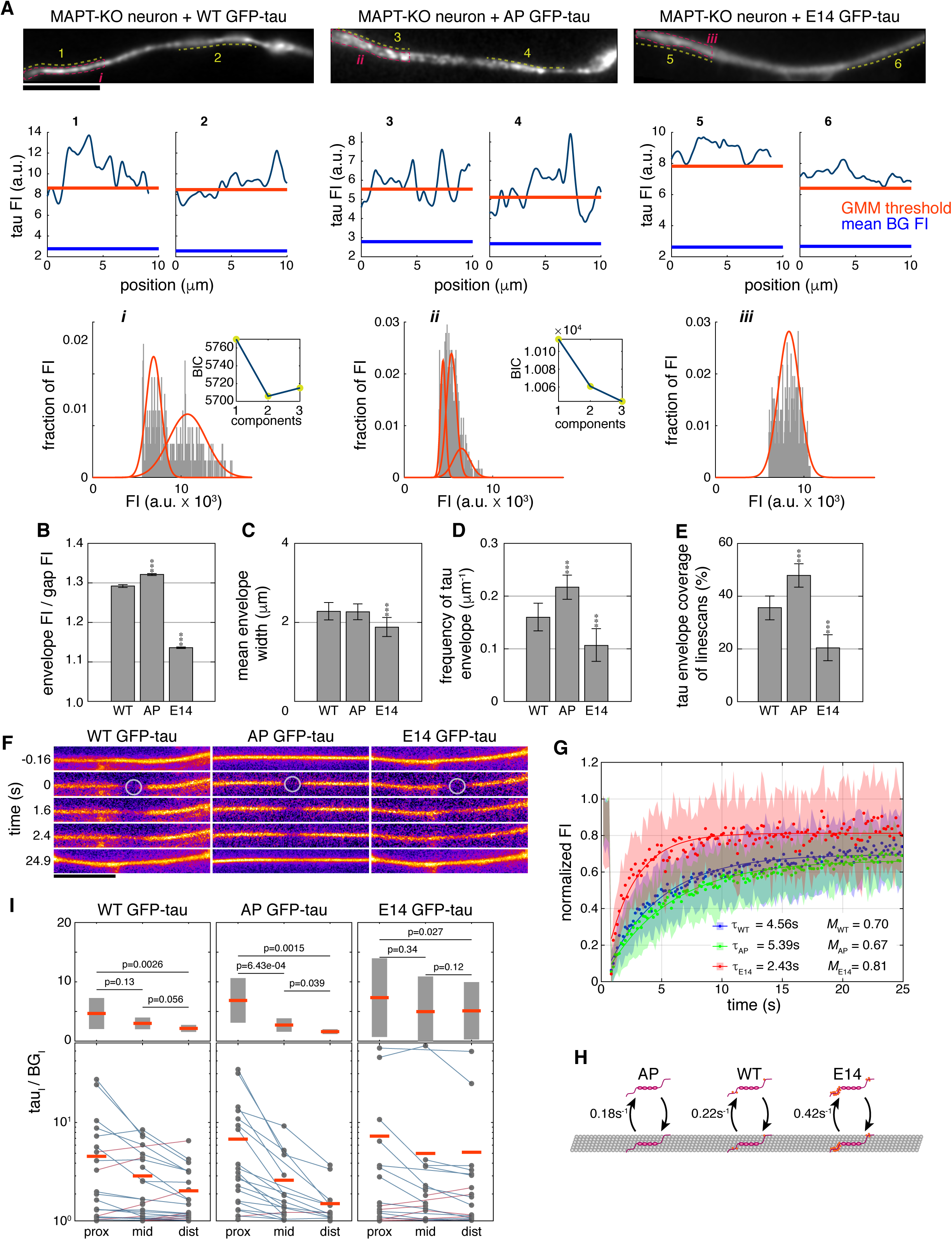
Tau hyperphosphorylation reduces the formation of envelopes in live neurons. **A)** Representative images of axons with tau envelopes present in iPSC-derived MAPT-KO neurons expressing WT (*n*=39), AP (*n*=39), or E14 (*n*=28) GFP-tau (4R0N). Each image is a max projection of 10 frames from live timelapse imaging. Images are all oriented so that the soma is towards the left and the distal axon is towards the right. Below, corresponding tau intensity plots from discrete locations (labeled 1–6) are shown. Horizontal lines indicating the GMM threshold (orange) and mean background fluorescence intensity (blue). Histograms of tau fluorescence intensity, fitted with a GMM to define the envelope threshold, are displayed for selected areas (labeled i–iii) marked by magenta ROIs. Insets show BIC analyses for multimodal distributions. **B–C)** Bar plots comparing B) tau envelope intensity and C) the mean envelope width of WT, AP, and E14 tau. **D–E)** Bar plots quantifying tau envelope frequency within discrete locations (D) and the percentage of axons covered by tau envelopes (E). Error bars indicate 95% CI. **F)** Timelapse images of fluorescence recovery after photobleaching assays of MAPT-KO neurons expressing WT(*n*=20), AP(*n*=21), or E14 (*n*=20) GFP-tau. Circles indicate the bleached regions. Note that tau envelopes are not clearly evident due to shorter exposure times needed for FRAP experiments and due to selection of higher GFP-tau expressing cells to ensure measurable recovery dynamics. **G)** Fluorescence recovery curves of WT (blue), AP (green), and E14 tau (red). Shaded regions indicate SD. The characteristic recovery (τ) and mobile fraction (*M*) are indicated on the plot for each tau construct. **H)** Schematic illustrates how tau phosphorylation influences its dissociation from microtubules. Hyperphosphorylated E14 tau dissociates more readily than WT tau or the phospho-resistant AP tau, which remain more stably bound (Fig S2). **I)** Plots show how the ratio of tau signal in axons of neurons expressing WT (*n*= 37), AP (*n*=32), and E14 (*n*=36) tau over background intensity varies for each tau construct across the proximal, mid, and distal axonal regions. In the bottom plot, blue lines indicate a decrease in the signal from the proximal towards the distal axon, and red lines indicate an increase in tau signal towards the distal axon. The top plot shows the means and grey bars represent SD. Orange bars show means. Wilcoxon signed rank test was used to determine the pairwise comparison of tau intensity between each axonal region as shown and the p-values are indicated in the above inset. The scale bars are 10 µm (A) and 5 µm (F). (*p < 0.05, **p < 0.001, ***p < 0.0001).

To further characterize the effects of tau hyperphosphorylation in cells, we performed fluorescence recovery after photobleaching (FRAP) assays to measure tau kinetics in axons. MAPT-KO neurons expressing WT, AP, or E14 GFP-tau were photobleached in a diffraction-limited spot, and fluorescence recovery was monitored (Fig 2F). AP tau recovered more slowly than WT tau, while E14 tau recovered the fastest (WTτ: 4.56s; APτ: 5.39s; E14τ: 2.43s) (Fig 2G). The 1D geometry of axons constrains diffusion, so we repeated FRAP in COS-7 cells, which have a flatter, more 2D-like geometry. Recovery was ∼2× faster in COS-7 cells (Fig S2).

Furthermore, in COS-7 cells, WT and AP tau showed clear microtubule enrichment as the fluorescence signal along microtubules recovered above cytosolic levels, while E14 tau showed no microtubule enrichment, consistent with our in vitro data showing reduced microtubule enrichment (Fig S1C). Thus, faster recovery rates would be expected if hyperphosphorylated tau was less stably bound to microtubules (Fig 2H).

Tau’s affinity for microtubules might also affect its distribution along the axon. We performed a pair-wise comparison of the mean ratio of tau intensity over background to test how phosphorylation impacts the distribution of tau across the proximal, mid, and distal axonal regions. We found that WT and AP tau are more enriched proximally than E14 tau, which is more uniformly distributed along the axon (Fig 2I). Altogether, these results suggest that hyperphosphorylation disrupts tau’s cooperative microtubule binding in neurons, preventing the formation of envelopes along microtubules, increasing exchange between microtubules and the cytosol, and altering tau distribution throughout axons (Fig 2I). Thus, hyperphosphorylation may weaken tau’s ability to regulate motor engagement and thereby contribute to axonal transport defects.

### Tau phosphorylation differentially regulates kinesin-1 and kinesin-3 motility along microtubules

We next sought to determine how tau phosphorylation influences the motility of motor proteins involved in axonal transport. Kinesin-1 and kinesin-3 are major drivers of anterograde transport in axons, trafficking endosomes, lysosomes, mRNA, and other cargoes (Jenkins et al., 2012; Beaudet et al., 2024; Nagpal et al., 2024). Previous studies showed that tau differentially regulates motor proteins, with kinesin-1 and kinesin-3 being more sensitive to tau-mediated inhibition, than kinesin-2 and dynein (Dixit et al., 2008; Vershinin et al., 2007; McVicker et al., 2011; Hoeprich et al., 2014; McVicker et al., 2014; Siahaan et al., 2019; Tan et al., 2019; Siahaan et al., 2022). However, how phosphorylation influences tau’s regulation of motors, particularly those most sensitive to tau, remained poorly understood. To address this question, we used TIRF-based reconstitution assays to examine how tau phosphorylation influences the behaviour of kinesin-1 and kinesin-3. Halo-tagged KIF5C (kinesin-1) and KIF1A (kinesin-3) motors expressed in COS-7 cells (Budaitis et al., 2022), were reconstituted along immobilized taxol-stabilized microtubules in the presence of extracts from mock-transfected cells or cells expressing WT, AP, or E14 GFP-tau. The fraction of processive KIF5C motility was slightly reduced in the presence of tau compared to mock extracts without tau (mock: 0.79 ± 0.05; WT: 0.71 ± 0.06; AP: 0.72 ± 0.05: E14: 0.72 ± 0.04) (Fig 3A, B). In addition, tau reduced the mean run lengths and velocities relative to mock conditions but with minimal differences among the phospho-variants ([mock: 1.91 ± 0.10 μm; WT: 1.59 ± 0.12 μm; AP: 1.51 ± 0.09 μm; E14: 1.62 ± 0.09 μm], [mock: 0.94 ± 0.04 μm/s; WT: 0.81 ± 0.04 μm/s; AP: 0.88 ± 0.03 μm/s; E14: 0.83 ± 0.04 μm/s]) (Fig 3E, S3A and C). In contrast, tau significantly reduced the number of KIF1A motors that engaged along microtubules (Fig 3C). For those KIF1A motors that did engage microtubules, the frequency of processive motility was significantly reduced among phospho-variants (mock: 0.85 ± 0.05; WT: 0.43 ± 0.13; AP: 0.14 ± 0.04; E14: 0.24 ± 0.07) (Fig. 3C, D). KIF1A mean run lengths were reduced substantially compared to mock conditions, with E14 tau having the strongest reduction, compared to AP and WT tau (mock: 3.48 ± 0.20 μm; WT: 2.30 ± 0.71μm; AP: 1.96 ± 1.06μm; E14: 1.66 ± 0.04μm), but the mean velocities remained largely unchanged (mock: 1.67 ± 0.05 μm/s; WT: 1.42 ± 0.41 μm/s; AP: 1.38 ± 0.41 μm/s; E14: 1.49 ± 0.38 μm/s) (Fig. 3F, S3D and F).

**Figure 3.**
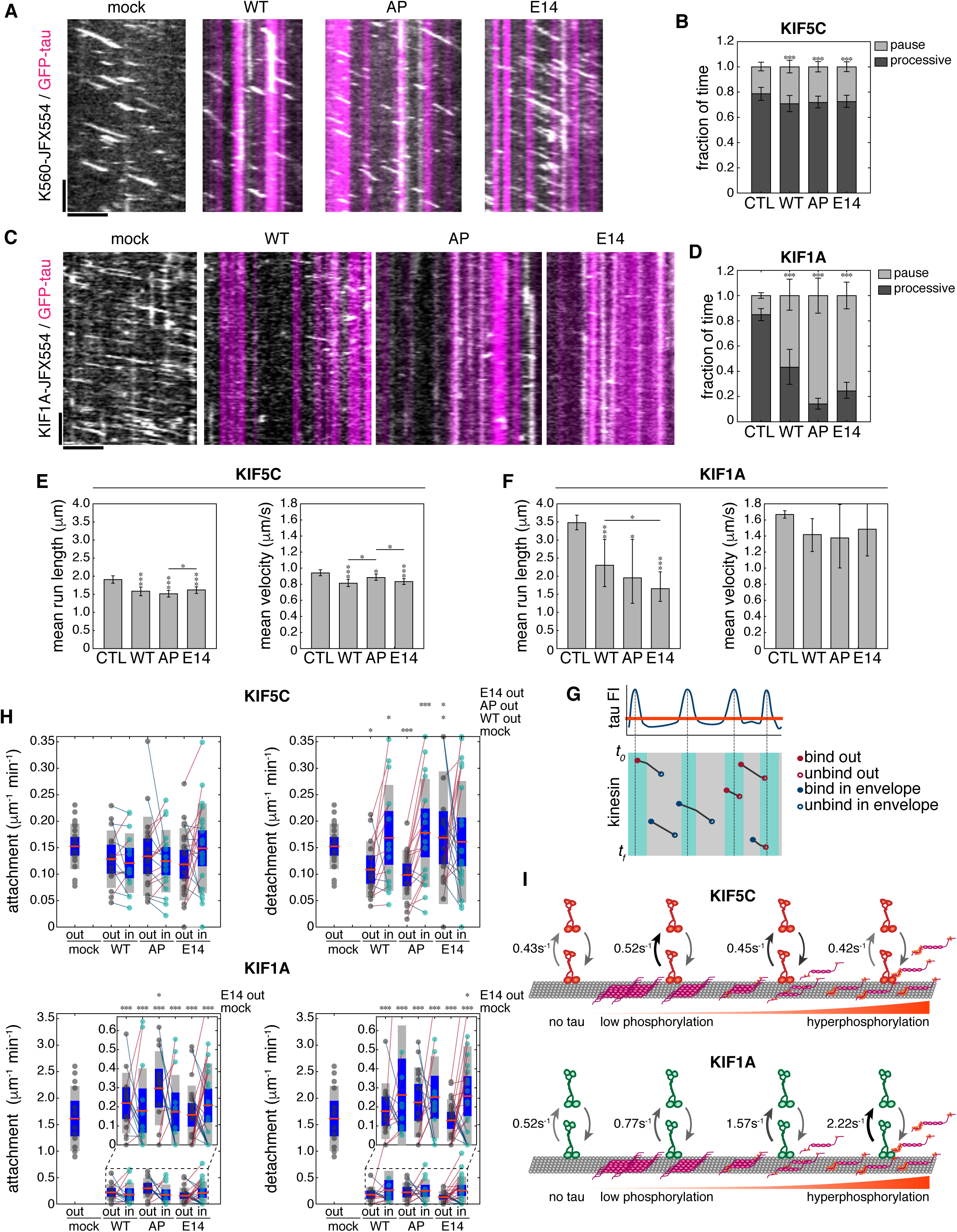
Tau phosphorylation differentially regulates kinesin-1 and kinesin-3 motility. **A and C)** Kymographs of constitutively active Janelia Fluor 554 (JFX554)- labelled A) KIF5C(1–560)-JFX554 and C) KIF1A(1–393)-JFX554 motors moving along taxol-stabilized microtubules incubated with mock cell lysate or lysates containing 500 nM WT, AP, or E14 GFP-tau (magenta). KIF5C (mock: *n*=1016, WT: *n*=792, AP: *n*= 1078, E14: *n*=1203) KIF1A (mock: *n*= 864, WT: *n*=173, AP: *n*=218, E14: *n*=305) over 3–4 replicates. **B and D)** Bar graphs show the fraction of time that kinesin motors are paused or exhibit processive motility. Error bars indicate 95% CI. **E–F)** Bar graph shows the bootstrapped mean run length and mean velocity of KIF5C (E) and KIF1A (F) ± WT, AP, E14 GFP-tau. Error bars indicate SEM. **G)** Schematic illustrating the approach used to assess the impact of tau on kinesin-microtubule attachment and detachment. Motors were observed attaching to or detaching from microtubules either within tau envelopes or outside of them. **H)** Paired sample plots show the attachment and detachment frequencies of KIF5C (top) and KIF1A (bottom). Red lines indicate increased frequency inside envelopes compared to outside and blue lines indicate decreased frequencies inside of envelopes. Insets for KIF1A show a zoomed-in view of the attachment and detachment frequencies. Blue bars indicate 95% CI, orange lines mark mean values, and grey bars denote SD. **I)** Schematic illustrating the impact of tau on the detachment kinetics of kinesin motors. KIF5C is weakly inhibited by WT and phospho-resistive tau compared to hyperphosphorylated tau that has a similar dissociation rate compared to KIF5C in mock conditions. Conversely, KIF1A is more strongly inhibited by tau hyperphosphorylation and dissociates at a faster rate compared to WT and AP tau. (*p < 0.05, **p < 0.001, ***p < 0.0001). Horizontal scale bars are 5 µm, vertical scale bars are 5 sec.

We next examined localized kinesin behaviour outside versus inside of tau envelopes by measuring attachment and detachment frequencies. Tau envelopes were identified using the GMM method described above (Fig. 1C), and each kinesin trajectory was classified based on whether it started or ended inside or outside of an envelope (Fig. 3G). Although E14 tau did not form envelopes like WT and AP tau, we instead measured attachment and detachment events defined by regions of high or low tau intensity determined using the GMM threshold. The attachment frequency of KIF5C was equally probable inside and outside of envelopes for all conditions ([mock: 1.52 ± 0.17 μm^−1^ min^−1^], [WT_out_: 1.28 ± 0.27 μm^−1^ min^−1^ vs WT_in_: 1.21 ± 0.28 μm^−1^ min^−1^], [AP_out_: 1.34 ± 0.34 μm^−1^ min^−1^ vs AP_in_: 1.24 ± 0.26 μm^−1^ min^−1^], [E14_out_: 1.18 ± 0.27 μm^−1^ min^−1^ vs E14_in_: 1.49 ± 0.34 μm^−1^ min^−1^]) (Fig. 3H). However, KIF5C detached more frequently inside of WT and AP tau envelopes than outside, whereas the detachment frequency was similar inside and outside of regions of increased E14 tau intensity ([mock: 1.52 ± 0.17 μm^−1^ min^−1^], [WT_out_: 1.09 ± 0.27 μm^−1^ min^−1^ vs WT_in_: 1.69 ± 0.51 μm^−1^ min^−1^], [AP_out_: 0.99 ± 0.21 μm^−1^ min^−1^ vs AP_in_: 1.78 ± 0.46 μm^−1^ min^−1^], [E14_out_: 1.69 ± 0.50 μm^−1^ min^−1^ vs E14_in_: 1.61 ± 0.46 μm^−1^ min^−1^]) (Fig. 3H). In contrast, KIF1A attachment frequency was significantly reduced whether inside or outside of envelopes compared to mock conditions ([mock: 1.61 ± 0.34 μm^−1^ min^−1^], [WT_out_: 0.22 ± 0.08 μm^−1^ min^−1^ vs WT_in_: 0.18 ± 0.12 μm^−1^ min^−1^]), [AP_out_: 0.30 ± 0.10 μm^−1^ min^−1^ vs AP_in_: 0.17 ± 0.10 μm^−1^ min^−1^], [E14_out_: 0.16 ± 0.06 μm^−1^ min^−1^ vs E14_in_: 0.21 ± 0.09 μm^−1^ min^−1^]). With WT and AP tau, KIF1A detachment frequency was not significantly different outside vs inside envelopes, but detached more frequently in areas of increased E14 tau intensity ([mock: 1.61 ± 0.34 μm^−1^ min^−1^], [WT_out_: 0.18 ± 0.08 μm^−1^ min^−1^ vs WT_in_: 0.26 ± 0.19 μm^−1^ min^−1^], [AP_out_: 0.22 ± 0.10 μm^−1^ min^−1^ vs AP_in_: 0.25 ± 0.11 μm^−1^ min^−1^], [E14_out_: 0.13 ± 0.11 μm^−1^ min^−1^ vs E14_in_: 0.26 ± 0.10 μm^−1^ min^−1^]), suggesting that KIF1A is more inhibited by diffusely bound, non-cooperative tau on microtubules (Fig. 3H).

Dissociation rate constants estimated from KIF5C and KIF1A dwell times also revealed significant differences in the presence of WT, AP, and E14 tau and mock conditions (Fig. 3I). KIF5C dissociation rates were similar on tau-free microtubules and in the presence of E14 tau, but were slightly higher with WT tau and AP tau (mock: 0.43 s^−1^; WT: 0.45 s^−1^; AP: 0.52 s^−1^; E14: 0.42 s^−1^) (Fig. 3I, S3B). In contrast, the dissociation rate of KIF1A increased substantially in the presence of tau, with E14 tau having the highest increase followed by WT, then AP tau (mock: 0.52 s^−1^; AP: 0.77 s^−1^; WT: 1.57 s^−1^; E14: 2.22 s^−1^) (Fig. 3I, S3E). Because E14 tau exhibited lower intensity on microtubules compared to WT and AP tau, we asked whether the effects on kinesin motility varied as a function of local tau intensity. To address this, we performed Spearman’s correlation analysis comparing tau pixel intensities along microtubules with standard deviation projections of kinesin intensity, where brighter pixels indicate higher dynamics. Across all tau phospho-variant, kinesin dynamics were consistently higher in regions with lower tau intensity (Fig S3G and H). However, KIF5C remained more dynamic in regions of elevated E14 tau intensity than at comparable WT or AP tau intensities (Fig S3G). In contrast, KIF1A showed greater dynamics in regions of high AP tau intensity than at similar E14 or WT tau intensities (Fig S3H). Together, these results indicate that kinesin dynamics are generally reduced in regions of high tau occupancy, but that KIF5C is less inhibited by E14 tau, whereas KIF1A is less inhibited by AP tau, at comparable local intensities. Because GFP-tau containing cell lysates were used to examine the effects of hyperphosphorylation on kinesin motility, endogenous MAPs and motors present in the lysates could potentially contribute to the observed effects. However, any such contributions would be expected to be similar across phospho-variants. Therefore, observed differences in motility likely reflect changes in tau phosphorylation state rather than variability arising from residual lysate components. These findings support the model that tau acts as a selective barrier to motor proteins, influencing cargo transport based on motor sensitivity. The differential inhibition of kinesin-1 and kinesin-3 suggests tau may gate transport by selectively modulating specific motors. Importantly, tau hyperphosphorylation appears to weaken its inhibitory effect on kinesin-1 while maintaining or enhancing its suppression of kinesin-3, potentially contributing to cargo transport deficits in neurodegenerative disease.

### Hyperphosphorylation alters directed lysosome transport and distribution in neurons

Axonal cargoes are transported by teams of motors with differing sensitivities to tau, as we observed for kinesin-1 (KIF5C) and kinesin-3 (KIF1A) (Fig 3) and as shown previously (Dixit et al., 2008; Vershinin et al., 2007; McVicker et al., 2011; Hoeprich et al., 2014; McVicker et al., 2014; Siahaan et al., 2019; Tan et al., 2019; Siahaan et al., 2022). We therefore asked how tau hyperphosphorylation influences the transport of cargoes driven by multiple motors. To test this, we examined lysosome transport. Like other endocytic vesicles, lysosomes are transported by teams of kinesin-1, -2, and -3 and dynein motors (Jordens et al., 2001; Brown et al., 2005; Loubéry et al., 2008; Rosa-Ferreira and Munro, 2011; Bentley et al., 2015; Pu et al., 2016; Beaudet et al., 2024). Lysosomes were labelled with LysoTracker, and their motility was monitored along axons (∼50 µm from either the soma or growth cone) in control, MAPT-KO, and MAPT-KO neurons expressing WT, AP, or E14 GFP-tau at DIV 7–8 (Fig. 4A). Kymographs were generated from timelapse image sequences, and lysosome trajectories were identified using KymoButler and analyzed by custom scripts.

**Figure 4.**
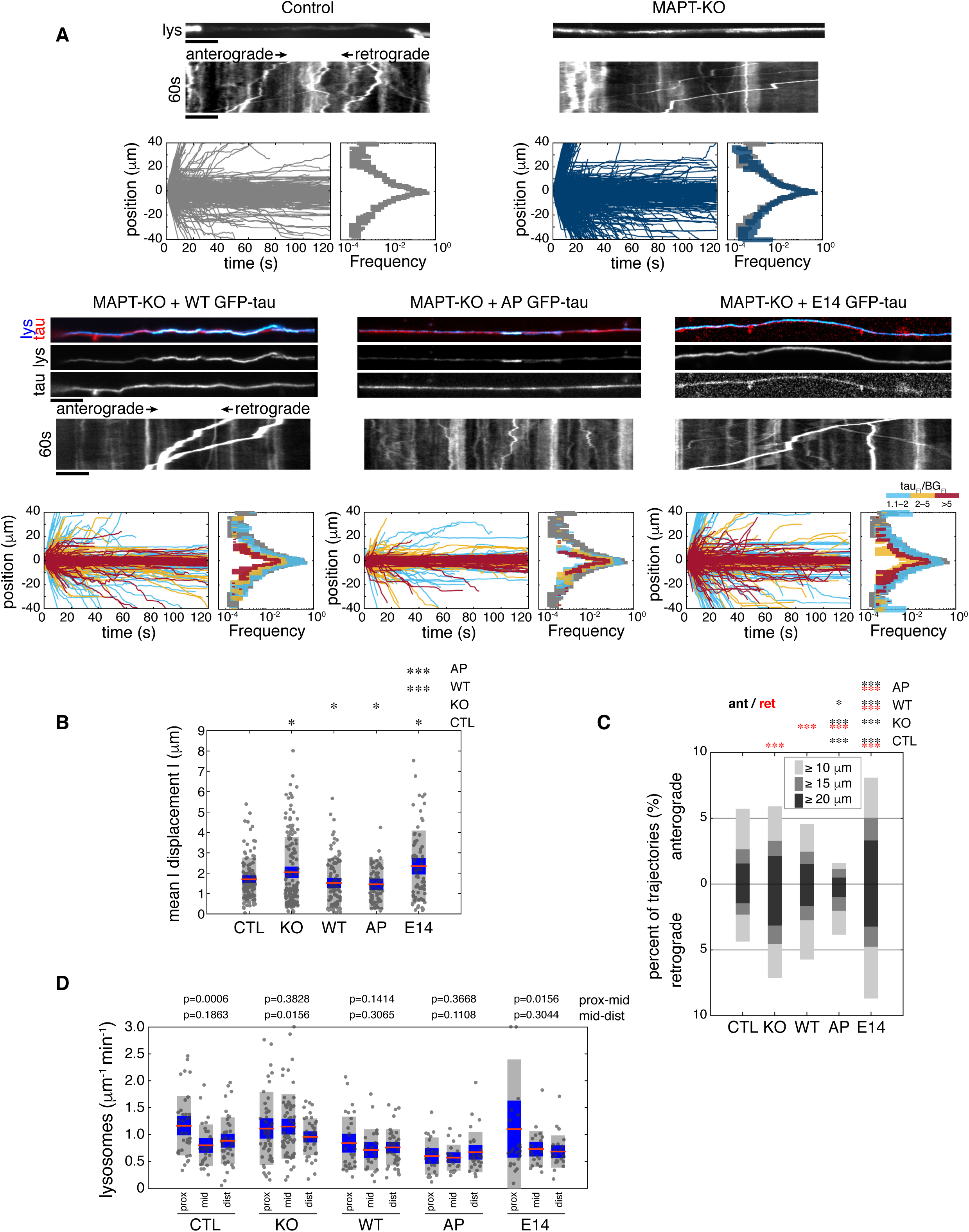
Tau perturbations impact the trafficking of lysosomes in neurons. **A)** Images show mid-axonal GFP-tau signal and max projections of lysosomes (lys) in control, MAPT-KO, and MAPT-KO neurons expressing WT, AP, and E14 GFP-tau. Below, kymographs show anterograde and retrograde lysosome transport for each condition. The number of cells analyzed for each condition are indicated in Table 2. Below each set of images, plots show the total number of lysosome trajectories mapped as position over time. Histograms next to each plot show the frequency distribution of lysosome travel distances for each condition. Positive values indicate anterograde transport (towards distal axon), while negative values indicate retrograde transport (towards soma). Trajectories are color-coded based on GFP-tau expression level: light blue (1.1–2× background), orange (2–5×), and red (>5×). Corresponding travel distance histograms are color-coded and overlaid with control data (grey) for comparison. **B)** Plots show the mean absolute displacements for each condition. **C)** Bar plot shows the fraction of anterograde and retrograde long-distance trajectories for control (CTL), MAPT-KO (KO), and MAPT-KO neurons expressing WT, AP, or E14 GFP-tau. Above the plot, asterisks represent statistical significance tested for anterograde (black) and retrograde (red) trajectories (* p < 0.05, ** p < 0.001, *** p < 0.0001)**. D)** Plots show the frequency of lysosomes along axons in the proximal, mid, and distal axon for each condition. Proximal regions were defined as ∼50 µm from the soma, mid-axonal regions as ∼halfway along the axon, and distal regions as ∼50 µm from the axon terminal. Statistical significance is indicated by asterisks above the at the top of plots (*p < 0.05, **p < 0.001, ***p < 0.0001). For B and D, Blue bars indicate 95% CI, orange lines mark mean values, and grey bars denote SD. Scale bars are 10 µm.

During each 2-minute timelapse sequence, lysosomes were predominantly immotile or exhibited short, diffusive trajectories, while only a small fraction of total transport was directed (Fig 4A). Mean displacements in WT- and AP-expressing MAPT-KO neurons were similar to controls (CTL: 1.70 ± 0.20 µm; WT:1.52 ± 0.24 µm; AP:1.46 ± 0.28 µm), whereas E14 tau expression increased mean displacements, similar to MAPT-KO neurons (MAPT-KO: 2.05 ± 0.27 µm vs E14: 2.34 ± 0.40 µm) (Fig 4B). We next examined the effects of tau on the fraction of long-range directed transport. The fraction of long-range retrograde transport >15 µm significantly increased in MAPT-KO neurons compared to controls (KO_ret_: 4.51% vs CTL_ret_: 2.57%), whereas the fraction of anterograde directed long-range transport events increased marginally (KO_ant_: 3.22% vs CTL_ant_: 2.26%) (Fig 4A and C). The fraction of long-range transport in WT tau-expressing MAPT-KO neurons was similar to controls (WT_ant_: 2.39%; WT_ret_: 2.69%). However, AP tau more significantly reduced long-range anterograde transport than retrograde transport (AP_ant_: 1.08% vs AP_ret_: 1.97%), whereas E14 tau increased long-range transport in both directions (E14_ant_: 4.97%; E14_ret_: 4.70%) (Fig 4C). Similar trends were observed when long-range lysosome transport was defined using thresholds of >10 µm or >20 µm (Fig 4C). Thus, indicating that phospho-resistive tau is inhibitory to long-range anterograde directed transport, while hyperphosphorylation, similar to tau-knockout, reduces inhibition of long-range transport in both directions.

We also assessed if the effects of tau phosphorylation on lysosome motility varied across different axonal regions and as a function of tau expression level (Fig 4A, S4A). We quantified the fraction of long-range lysosome transport in proximal (∼50 μm from the soma, avoiding the axon initial segment), mid (∼midpoint), and distal (∼50 μm from the growth cone) axonal regions (Fig S4A). Expression levels were measured as the ratio of mean GFP-tau pixel intensity within the axon relative to the mean background pixel intensity. While some variability was observed between each region, the effects on long-range lysosome transport were primarily dependent on tau expression level and scaled differently with each phospho-variant. With WT tau expression, lysosomes underwent long-range transport less frequently at high expression levels (>5× background signal) compared to neurons with low (1.1–2.0×) or intermediate (2.0–5.0×) expression (Fig 4A, S4B and C). With AP tau, the reduction in long-range transport was more pronounced with lower expression levels, and higher expression more strongly reduced anterograde directed transport (Fig 4A, S4B and C). Low expression of E14 tau increased long-range lysosome transport but was reduced at high expression levels, although to a lesser extent than in neurons expressing WT or AP tau (Fig. 4A, S4B and C).

One possible consequence of altered long-range transport is a change in lysosome distribution along the axon, which would be expected to impact degradative pathways and proteostasis. Therefore, we quantified how tau phosphorylation affects the number of lysosomes in proximal, mid, and distal axonal regions (Fig 4D). In control neurons, lysosomes were slightly enriched in the proximal axon relative to mid and distal regions, whereas in MAPT-KO neurons, lysosomes were distributed more uniformly across regions. In MAPT-KO neurons expressing WT tau, lysosome frequency was reduced compared to control neurons, but were marginally enriched in the proximal axon. In contrast, AP tau expression resulted in fewer lysosomes that were distributed more evenly across axonal regions, whereas E14 tau expression led to greater lysosome enrichment in the proximal axon (Fig 4D). Together, these data suggest that lysosomes are normally slightly enriched in the proximal axon and that tau promotes lysosome flux toward this region. AP-tau, which predominately binds cooperatively, blocks the proximal enrichment of lysosomes. Tau misregulation might disrupt lysosome localization by enhancing long-range retrograde transport, and therefore degradative pathways. These effects would likely become more pronounced over longer timescales and in more mature or aged neurons.

Hyperphosphorylated tau enhances retrograde processive lysosome motility, while phospho-resistant tau strongly inhibits anterograde processive motility

Lastly, we asked which aspects of lysosome transport are most affected by tau hyperphosphorylation that would disrupt long-range transport and lysosome distribution. Lysosomes are bidirectional organelles that frequently reverse directions and alternate between stationary, diffusive, and processive phases of motility. We therefore examined how each tau phospho-variant affected transport directionality and the different phases of motility. Trajectories were segmented by identifying directional reversal events, and the intervals between reversals were classified based on displacement as stationary (less than 0.16 µm, within the tracking uncertainty), diffusive (between 0.16 µm and 1.2 µm), or processive (greater than 1.2 µm) (Fig 5A). Lysosomes were predominantly stationary or diffusive, with processive motility accounting for only a small fraction of total movement (Fig 5B). The fraction of processive motility in MAPT-KO neurons was similar to controls (CTL: 7.37 ± 0.55% vs MAPT-KO: 6.76 ± 0.51%), although lysosomes underwent more frequent directional reversals (CTL: 0.83 ± 0.04 μm^−1^ min^−1^ vs KO: 0.92 ± 0.04 μm^−1^ min^−1^) (Fig 5B and C). The fraction of processive motility with AP tau was similar to WT tau (AP: 5.60 ± 0.56% vs. WT: 5.93 ± 0.59%) but reduced the frequency of directional reversals (AP: 0.70 ± 0.053 μm^−1^min^−1^ vs. WT: 0.83 ± 0.05 μm^−1^ min^−1^). In contrast, E14 tau increased the fraction of processive motility (7.92 ± 0.72%) while reversal frequencies were similar to WT (0.88 ± 0.07 μm^−1^min^−1^) (Fig 5B and C).

**Figure 5.**
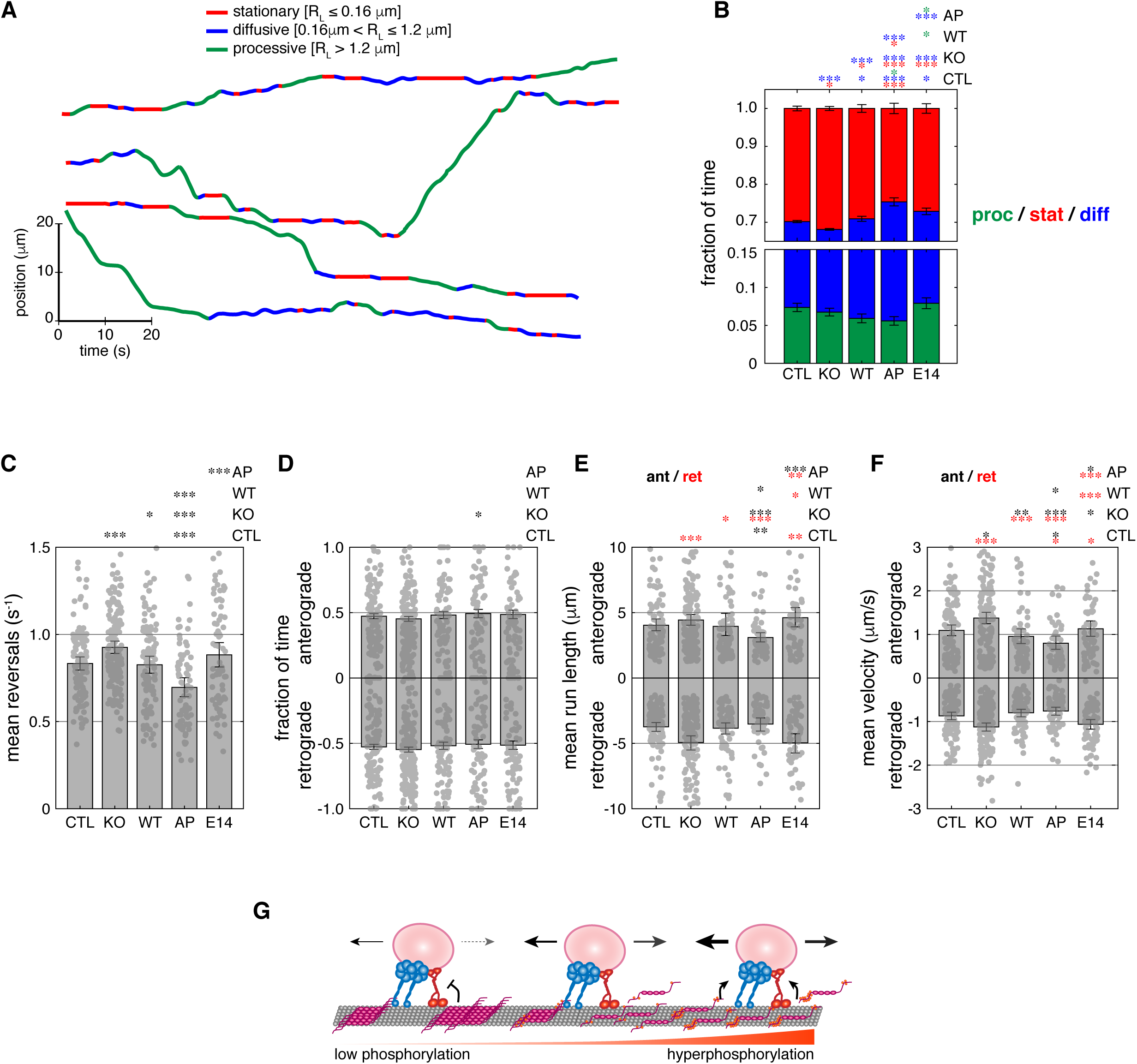
Tau hyperphosphorylation relieves inhibition of processive lysosome motility in neurons. **A)** Shown are examples of lysosome trajectories segmented into stationary (red), diffusive (blue), and processive (green) motility based on run length. Stationary segments were defined as periods between two reversal events with a run length (R_L_) < 0.16 µm; diffusive segments as those with run lengths between 0.16 µm and 1.2 µm; and processive segments as those > 1.2 µm. **B)** Plots show the fraction of time lysosomes exhibited processive, diffusive or stationary phases of motility for control (CTL), MAPT-KO (KO), and MAPT-KO neurons expressing WT, AP, and E14 GFP-tau. Error bars represent 95% CI. Statistical significance is indicated above the plots for comparison of the fraction of processive (proc; green) diffusive (diff; blue), and stationary (stat; red) time for each condition. **C–D)** Bar plots show C) the mean reversal frequency of lysosomes, D) and the fraction of time of anterograde or retrograde directed processive motility. **E–F)** Bar plots show E) the mean run lengths and F) mean velocities of processive runs of lysosomes for each condition. Error bars in C), E), and F) indicate SEM and in D) indicate 95% CI. The number of trajectories and cells analyzed under each condition are indicated in Table 2. **G)** Schematic summarizing the impact of tau hyperphosphorylation on bidirectional lysosome transport in neurons. Low tau phosphorylation reduces processive anterograde transport but has no significant impact on retrograde transport, whereas hyperphosphorylated tau relieves inhibition of anterograde transport and enhances retrograde transport, similar to the effects observed in tau knockout conditions. Statistical significance is shown above each plot, where red asterisks indicate comparisons of retrograde transport and black asterisks indicate comparisons of anterograde transport. (* p < 0.05, ** p < 0.001, *** p < 0.0001).

Across all conditions, the fraction of anterograde and retrograde directed processive motility were similar (Fig 5D). However, the mean run lengths and velocities of processive runs differed significantly between conditions. In MAPT-KO neurons, retrograde-directed lysosomes were more processive than in controls, exhibiting increased mean run lengths (CTL_ret_: 3.74 ± 0.34 μm vs KO_ret_: 4.94 ± 0.55 μm) and velocities (CTL_ret_: 0.87 ± 0.09 μm/s vs KO_ret_: 1.12 ± 0.09 μm/s) (Fig 5E and F). Lysosome processivity in MAPT-KO neurons expressing WT tau was similar to controls. However, AP tau reduced anterograde processive transport, as reflected by shorter run lengths (AP_ant_: 3.10 ± 0.35 μm vs WT_ant_: 3.93 ± 0.83 μm) and lower velocities (AP_ant_: 0.80 ± 0.15 μm/s vs WT_ant_: 0.95 ± 0.17 μm/s). By comparison, E14 tau expression caused a marginal increase in anterograde processive run lengths and velocities, while significantly enhancing retrograde processivity, demonstrated by increased mean run lengths (E14_ret_: 4.97 ± 0.70 μm vs WT_ret_: 3.84 ± 0.41 μm/s) and velocities (E14_ret_: 1.06 ± 0.11 μm/s vs WT_ret_: 0.80 ± 0.09 μm/s). Together, these results indicate that tau phosphorylation state differentially regulates processive lysosome motility. AP tau reduced directional switching and is more inhibitory to anterograde-directed processive motility, whereas E14 tau alleviated this inhibition and enhanced retrograde processivity, resembling tau knock-out conditions. Overall, these finding demonstrate that changes in tau phosphorylation alter the balance of directed lysosome transport, leading to altered lysosome flux and distribution along axons.

## Discussion

Tau hyperphosphorylation is a hallmark of Alzheimer’s Disease and associated with the aggregation of tau into neurofibrillary tangles. Here, we examined the functional consequences of hyperphosphorylation on tau’s roles in organizing the cytoskeleton and regulating intracellular transport. We find that hyperphosphorylation disrupts tau’s ability to form envelopes on microtubules and alters microtubule-based transport. Tau binds microtubules cooperatively to form cohesive envelopes in live neurons, consistent with its behaviour on stabilized microtubules in vitro (Figs 1 and 2). Disease-related hyperphosphorylation prevents cooperative binding and enhances tau’s dissociation from microtubules (Figs 1 and 2). Tau hyperphosphorylation also has different effects on different kinesin motors that transport several types of axonal cargoes. KIF5C motility is mildly inhibited by mammalian expressed tau compared to KIF1A that is strongly inhibited (Fig 3). Hyperphosphorylation reduces KIF5C dissociation from microtubules, while it increases that of KIF1A (Fig 3H), suggesting that misregulation of tau could alter axonal trafficking by disrupting kinesin activity. In agreement, we found that hyperphosphorylation weakens tau-mediated inhibition of lysosome transport in neurons. While unphosphorylated tau acts as a strong barrier to anterograde lysosome movement, hyperphosphorylated tau alleviates this inhibition and enhances retrograde directed transport, mimicking tau knockout conditions in which lysosomes moved more processively in both directions (Figs 4 and 5). These disruptions in transport, which result in altered lysosome flux along the axon (Fig 4D), likely lead to impairments of degradative pathways and protein homeostasis, which contribute to neuronal dysfunction and degeneration in tauopathies.

Isolating the effects of tau phosphorylation presents significant challenges. Tau is phosphorylated at numerous residues and at different developmental stages, and individual tau molecules often exhibit heterogeneous phosphorylation patterns (Lindwall and Cole, 1984; Mandelkow et al., 1995; Trinczek et al., 1995; Hasegawa et al., 1998; Siahaan et al., 2026 Fan et al., 2025). There are six tau isoforms expressed in the mature human brain (Weingarten et al., 1975; Goedert et al., 1988; Goedert et al., 1989a; Goedert et al., 1989b; Andreadis et al., 1992), further complicating efforts to define the contribution of specific phosphorylation events to its function. To isolate the effects of tau hyperphosphorylation, we employed phospho-mimetic and phospho-resistant tau variants (Hoover et al., 2010). To avoid effects from various endogenous tau isoforms, GFP-tau phospho-variants were expressed in COS-7 cells for in vitro reconstitution studies, where tau is not endogenously expressed, or in MAPT-KO neurons to assess transport phenotypes in the absence of endogenous tau. These approaches enabled us to examine tau function in a near-physiological environment, preserving relevant mammalian cytosolic factors and PTMs, while isolating the effects of hyperphosphorylation of a single tau isoform from the multiple isoforms of endogenous tau that are present in neurons. An additional consideration is that our study focused on a single tau isoform. iPSC-derived neurons at DIV 7–8 predominantly express 3R tau, with 4R tau expression increasing during maturation (Iovino et al., 2010; Iovino et al., 2015; Sposito et al., 2015). Since 3R tau has a lower affinity for microtubules but more strongly inhibits kinesin and dynein motors compared to 4R tau (Vershinin et al., 2008; McVicker et al., 2011; McVicker et al., 2014), hyperphosphorylation may have more severe effects in mature neurons where 3R and 4R tau are equally expressed. Future work is needed to explore how these isoforms interact, influence each other’s microtubule binding and cooperative assembly, and how phosphorylation heterogeneity among tau molecules (Siahaan et al., 2026; Fan et al., 2025) contributes to disease progression in tauopathies.

Our findings support the growing evidence that the proline-rich region and C-terminal pseudo-repeat domain of tau regulate its cooperative interactions with microtubules (Tan et al., 2019; Siahaan et al., 2019; Siahaan et al., 2022). The 14 disease-associated mutations used here to mimic hyperphosphorylation are located within these regions (Fulga et al., 2007; Steinhilb et al., 2007; Hoover et al., 2010), which together with the microtubule-binding repeats, comprise the minimal domains required for cooperative binding (Tan et al., 2019). Our results demonstrate that altering the phospho-state of the proline-rich region and pseudo-repeat domain directly affects how tau interacts with microtubules. Preventing phosphorylation strengthened cooperative binding, whereas phospho-mimetic mutations significantly reduced cooperative envelope formation and increased tau dissociation from microtubules, indicating weakened microtubule affinity (Figs 1, 2, S2). Other studies showed that additional sites within the N-terminal projection domain also contribute to tau cooperativity. For example, the pathogenic R5L mutation in the phosphatase-activating domain (PAD), disrupts cooperative binding by altering the structure of the N-terminal projection domain without affecting the microtubule-binding repeats or tau’s microtubule affinity (Cario et al., 2022). Likewise, mutations at Y18, the terminal residue of PAD, increase tau dynamics on microtubules, reduce tau accumulation, and impair tau-mediated regulation of intracellular transport (Kanaan et al., 2012; Stern et al., 2017; Balabanian et al., 2022). Together, these findings indicate that modifications within the N-terminal projection domain, either in PAD or the proline-rich region, as well as the C-terminal pseudo-repeat domain can alter tau-microtubule interactions and, consequently, tau’s ability to regulate intracellular transport. Several mechanisms could account for these effects. These domains may allosterically modulate the microtubule-binding repeats, phosphorylation may directly reduce microtubule affinity, or it may impair tau’s ability to compact the underlaying microtubule lattice (Siahaan et al., 2022), thereby affecting cooperative envelope formation and the interactions of other MAPs and motors with microtubules. Further studies will be required to distinguish between these possibilities and define the specific contributions of these regions to tau cooperativity and microtubule binding.

Previous single-molecule studies showed that kinesin-1 and kinesin-3 are more sensitive to inhibition by tau, whereas kinesin-2 and dynein are comparatively less affected. Consistent with tau’s proposed role as a selective barrier on microtubules, reducing tau-microtubule association through hyperphosphorylation or tau knockout, would be expected to enhance transport of cargoes driven by motors that are more sensitive to tau, such as kinesins-1 and -3. Our results demonstrate that tau phosphorylation differentially affects these motors. Kinesin-1 was more sensitive to phospho-resistant tau, showed by increased motor detachment within envelopes. In contrast, although kinesin-3 was strongly inhibited by tau overall (Fig 3C and D), hyperphosphorylated tau produced a greater inhibitory effect, as evidenced by accelerated motor detachment in regions of elevated E14 tau intensity along microtubules (Fig 3H). These findings indicate that phosphorylation alters tau’s regulation of kinesin-1 and kinesin-3 through distinct mechanisms. We examined KIF5C and KIF1A as representative members of the kinesin-1 and kinesin-3 families based on prior studies demonstrating their sensitivity to tau-mediated inhibition (Dixit et al., 2008; Vershinin et al., 2007; McVicker et al., 2011; Hoeprich et al., 2014; McVicker et al., 2014; Siahaan et al., 2019). However, the sensitivity of kinesin homologs to tau regulation likely varies due to differences in their intrinsic motor properties. Furthermore, endogenous cargo transport involves organelle-specific motor-adaptor complexes containing multiple redundant motor types and variable copy numbers (Beaudet et al., 2024). Thus, the effects on organelle trafficking, such as lysosome transport, are expected to be more nuanced than those observed for individual motors in vitro.

Lysosome motility is driven by multiple motor types, and the transport phenotypes observed here likely reflect the combined contributions of kinesins-1, -2, -3, and dynein. Phospho-resistant tau reduced anterograde transport with little effect on retrograde transport, whereas hyperphosphorylated tau relieved inhibition of anterograde transport and enhanced retrograde transport (Figs 4 and 5). Our results are consistent with a tug-of-war mechanism, in which tau acts as a selective barrier that preferentially inhibits kinesins-1 and -3, thereby shifting the balance towards dynein-driven transport (Müller et al., 2008; Chaudhary et al., 2018; Beaudet et al., 2024). Another possibility is that the teams of motors that transport cargo have distinct functions (Nagpal et al., 2024), including force-producing motors and others that act primarily as tethers (Feng et al.,2018; Arpag et al., 2019). In this context, enhanced detachment of kinesin-3 by hyperphosphorylated tau could reduce its tethering capacity, shifting the balance of forces in favor of dynein-driven retrograde transport. An alternative, but not mutually exclusive, explanation is that hyperphosphorylated tau alters tau occupancy along microtubules, allowing increased recruitment of other MAPs. For example, MAP7 competes with tau for lattice occupancy and has been shown to enhance kinesin-1 motility while inhibiting kinesin-3 (Monroy et al., 2018). These opposing effects could enhance anterograde transport through increased kinesin-1 processivity or promote retrograde transport by limiting kinesin-3 activity, while having a comparatively minor impact on dynein-mediated transport. Together, these findings suggest that the transport effects of tau hyperphosphorylation are determined not only by changes in tau-motor interactions but also by cargo-specific motor compositions and their differential sensitivities to tau (Beaudet et al., 2024). Overall, our results suggest that hyperphosphorylated tau fails to maintain its normal role as a regulatory barrier on microtubules. Future studies examining how tau phosphorylation influences the coordination of motors and other MAPs will be important for understanding how these molecular changes are integrated to regulate bidirectional transport in neurons.

Taken together, our findings show that tauopathy-related hyperphosphorylation weakens tau’s interaction with microtubules, reducing its ability to form cohesive envelopes and regulate axonal transport. Beyond impaired lysosome trafficking, this disruption likely affects signaling and other maintenance pathways essential for axonal health and function. Our results, which align with a recent study (Moretto et al., 2026), suggest a potential pathway, where prior to oligomerization and late-stage aggregation, hyperphosphorylated tau first impairs transport, which then contributes to axonal decline. As aggregation progresses, these effects are amplified, leading to further trafficking defects, axonal blockages, inflammation, and ultimately neurodegeneration. Overall, our results suggest that the disruption of axonal transport, caused by tau’s impaired association with microtubules, is one of the earliest drivers of neurodegeneration.

### Limitations of this study

This study combined in vitro reconstitution with iPSC-derived neurons to obtain mechanistic insight into how disease-related hyperphosphorylation alters tau’s cooperative association with microtubules and its function as a selective barrier for motor-driven cargoes. However, several limitations should be considered when interpreting these findings. The iPSC-derived neurons used here correspond to an early developmental stage and are more similar to embryonic or immature neurons than the aged neurons affected in Alzheimer’s disease and other tauopathies. Consequently, disease phenotypes in mature or patient-derived neurons may be more pronounced and involve additional regulatory mechanisms not captured in this system. This difference in disease stage may also explain discrepancies with previous studies. For example, Hallinan et al. (2019) reported lysosome transport defects with E14 tau opposite to those observed here, but their study used more mature hippocampal neurons and observed MC1-positive tau aggregates in axons. Thus, rather than being contradictory, the findings may reflect distinct stages of pathology, with our results capturing early effects of hyperphosphorylation on tau–microtubule interactions prior to aggregation, whereas later-stage oligomerization and aggregation may impair transport by clogging axons. Moreover, disease-associated phosphorylation is likely more heterogeneous and dynamic than the pseudo-phosphorylation mutants used here, and phosphorylation at different sites may differentially regulate tau function. In addition, DIV7–8 iPSC-derived neurons predominantly express 3R tau (Iovino et al., 2010; Iovino et al., 2015; Sposito et al., 2015), whereas our rescue experiments used 4R0N tau. Although lysosome transport in WT 4R tau-expressing MAPT-KO neurons was indifferent from controls, isoform-specific differences may influence the magnitude of phosphorylation-dependent effects, particularly in mature neurons where 3R and 4R tau isoforms are co-expressed. Finally, while COS-7 cells provide a robust mammalian expression system for tau and kinesins, they lack the full complement of neuronal regulatory factors and organization that shape MAP and motor behavior in neurons. Together, these limitations highlight that while our experimental systems enable precise mechanistic dissection of tau-microtubule and tau–motor interactions, future studies in more mature neuronal models and in patient-derived neurons will be essential to fully define how these mechanisms operate in the context of aging and neurodegenerative disease.

## Methods

### DNA constructs

The pRK5-EGFP-Tau (Addgene plasmid # 46904), pRK5-EGFP-Tau AP (Addgene plasmid # 46905), and pRK5-EGFP-Tau E14 (Addgene plasmid # 46904) plasmids were gifts from Karen Ashe. KIF1A(1-393)-LZ-HALO reported in (Budaitis et al., 2021) was generously provided by Kristen Verhey (U. Michigan). KIF5C(1-560)-HALO reported in (Twelvetrees et al., 2016) was generously provided by Erika Holzbaur (U. Pennsylvania).

### Cell culture and transfection

COS-7 monkey kidney fibroblast cells (ATCC) were maintained in DMEM (Gibco), supplemented with 10% fetal bovine serum (Thermo Fisher Scientific) and 1% GlutaMAX (Gibco) and cultured in MatTek glass-bottom dishes (No. 1.5 Coverslip) (MatTek Corporation, Ashland, MA) at 37°C with 5% CO_2_ for 24–48 h before transfection. Cells were then transfected with 500 ng of DNA plasmid prepared in OPTI-MEM Reduced Serum (Life Technologies), using Lipofectamine LTX with Plus-Reagent, according to the manufacturer’s instructions (Invitrogen, Thermo Fisher Scientific) for 24 h prior to imaging or homogenization.

Human neurons were induced from iPSC control and MAPT-KO AIW002-02 lines. The iPSCs were obtained from the C-BIG repository (Montreal Neurological Institute-Hospital, Montreal, QC, Canada) and were generated from the PBMC of a control 37-year-old male donor. MAPT knockout was achieved by CRISPR-Cas9 and verified by PCR and sequencing. Guide RNAs (gRNAs) were designed using “optimized CRISPR design” tool (www.crisp.mit.edu). The locations of gRNA1(GGATAAGTTCTGAGGAGTGT) and gRNA2 (TGATCTGGGCCTGCTGTGCA) were around exon 2 with ATG start codon. Oligonucleotides with Bsb1 cleavage overhang were ordered from Life Technologies, annealed and cloned into the PX459 Cas9/puromycin vector (Addgene plasmid #48139). iPSCs were characterized to ensure correct genotyping, pluripotency, and screened for microbiology/virus. iPSCs were maintained in mTeSR Plus media (STEMCELL Technologies, Vancouver, BC) on Matrigel (Corning, Corning, NY) coated culture dishes using Gentle Cell Dissociation Reagent (STEMCELL Technologies) for passaging. The iPSCs were cultured for ∼ 30 days in STEMdiff SMADi medium (STEMCELL Technologies) to induce the formation of neuronal progenitor cells (NPCs). NPCs were dissociated from culture dishes (Accutase Cell Dissociation Reagent) and seeded on poly-l-ornithine (PO) (10 µg/ml)/laminin (15 µg/ml)-coated MatTek glass-bottom dishes (No. 1.5 Coverslip) at 37°C with 5% CO_2_ in BrainPhys differentiation media (STEMCELL Technologies) at about 2,000 cells per cm^2^. Six days after seeding, the cells were transfected with lipofectamine LTX reagent at a 3:1 reagent:DNA according to the manufacturer’s instructions then imaged at 7–8 days following terminal differentiation induction.

### Live cell imaging

To capture lysosome dynamics, neurons were treated with 70 nM LysoTracker (Thermo Fisher Scientific, Waltham, MA) for 30 minutes at 37°C with 5% CO_2_ prior to replacing the media with BrainPhys containing 15 mM HEPES (H3375, Millipore Sigma). Live cell imaging was performed on an Eclipse Ti-E inverted microscope (Nikon) with custom optics for total internal reflection fluorescence (TIRF) and imaged using EMCCD camera (iXon U897; Andor Technology) maintained at 37°C. Time lapse recordings were acquired with 120 msec exposures using a 640 nm laser (100 mW) set at 1% power for 120 sec per cell using NIS-Elements acquisition software (Nikon). To image GFP-tau envelopes in neurons, imaging was performed as described but acquired with 500 msec exposures for 30 seconds using a 480nm laser set at 2% laser power. To measure GFP-tau expression in neurons a single frame was taken with 500nm exposure using a 480nm laser set at 2% laser power. Image files were exported as TIFFs, which were opened with ImageJ (NIH) and lysosomes were tracked by kymograph analysis.

Briefly, linescans were traced over the axon to generate kymographs, which were then analyzed using KymoButler (Jakobs et al., 2019) to identify individual lysosome trajectories. The coordinates of each trajectory were then imported into MATLAB (The MathWorks) to analyze the mode of transport, directionality, displacement, and number of reversals for all trajectories within cells using custom scripts.

### FRAP

COS-7 cells or neurons at 7–8 DIV expressing WT, AP, or E14 GFP-tau were used to assess tau kinetics in live cells. FRAP experiments were performed at 37 °C using a DeltaVision OMX system equipped with a 100×/1.40 NA objective lens and an EMCCD camera (Evolve, Photometrics). Prior to photobleaching, 5 pre-bleach images were acquired with 100 ms exposure. Photobleaching was performed within a diffraction limited spot using 50% 488 nm laser power for 50 ms. Fluorescence recovery was then recorded for 100 frames for COS-7 cells or 150 frames for neurons using 31% 488 nm laser power and 100 ms exposure per frame. Fluorescence recovery curves were generated using ImageJ, and background-corrected to account for acquisition-related photobleaching. Fluorescence recovery data were fit to a single-exponential function described by equation (1):

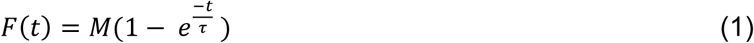

where *F*(*t*) is the fluorescence intensity at time *t*, *M* is the mobile fraction, *τ* is the characteristic recovery time and the dissociation rate constant (*k_off_*) is described by equation (1.1):

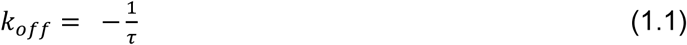

### COS-7 cell extract preparation

To prepare GFP-tau–containing cell extracts, COS-7 cells expressing WT, AP, or E14 GFP-tau were harvested 24 h post-transfection. For extracts containing kinesin motor proteins, COS-7 cells expressing Halo-tagged KIF5C or KIF1A were labeled with Janelia Fluor JFX554 ligand (Promega) using the manufacturer’s rapid labeling protocol and collected 24 h post-transfection. Cells were incubated with 200 nM JFX554 for 30 min, rinsed 2× with pre-warmed DMEM, followed by 2× PBS to remove residual media. Cells were then collected using a cell scraper in ice-cold lysis buffer (10 mM PIPES, 50 mM potassium acetate, 4 mM MgCl₂, 1 mM EGTA, pH 7.0), supplemented with protease inhibitor cocktail (BioShop), 10 mM DTT, and 0.5% Triton X-100. For motor-containing extracts, 1 mM MgATP was also included. Cell suspensions were incubated on ice for 1 min, then centrifuged at 1,000 × g for 10 min at 4 °C. Supernatants were transferred to fresh ice-cold microcentrifuge tubes and spun at 16,000 × g for 10 min at 4 °C. The concentration of GFP-tau in the extracts was determined by measuring absorbance at 488 nm using a Nanodrop spectrophotometer, applying an extinction coefficient of 56,000 M⁻¹cm⁻¹ (Table 1). The resulting soluble fractions containing GFP-tau or fluorescently labeled kinesin motors were aliquoted, flash-frozen in liquid nitrogen, and stored at –80 °C for subsequent experiments.

### Microtubule polymerization

To prepare taxol-stabilized microtubules, polymerization reaction mixtures containing 8% Alexa647-labeled tubulin and 92% unlabeled tubulin (final concentration of 5 mg/mL) in BRB80 (80 mM K-Pipes, 1 mM MgCl_2_, 1 mM EGTA, pH 6.8) supplemented with 1 mM GTP were prepared on ice thenn incubated for 25 mins at 37°C. Microtubules were then stabilized with 20 μM Taxol (Cytoskeleton) and incubated for 25 mins at 37°C. For GMPCPP-microtubules, reaction mixtures were mixed with 1mM GMPCPP (Jena Bioscience, Jena, Germany) for 60 minutes. Microtubules were cleared 2X by pelleting them at 10,600*g* for 5 mins at RT then washed with T-BRB80 (BRB80 supplemented with 20 μM Taxol).

### In vitro reconstitution assays

Flow chambers were first incubated with anti-β-tubulin (T4026 clone TUB2.1, Sigma) diluted 1:25 in BRB80 for 5 mins. Chambers were then treated with F-127 for 5 mins and washed 2X with T-BRB80. Fluorescently-labeled microtubules were added to the chambers and incubated for 5 mins at RT. Unbound microtubules were washed out with 2X T-BRB80. GFP-tau containing cell extracts were diluted 2-fold in MAB supplemented with 0.2 mg/ml BSA, 10 mM DTT, 20 mM Taxol, 15 mg/ml glucose, ≥ 2000 units/g glucose oxidase, ≥ 6 units/g catalase, and 1 mg/ml casein and flown through the chamber containing microtubules and imaged in TIRF. To image GFP-tau envelopes in vitro, images were acquired with 500 msec exposures for 30 seconds using a 480nm laser set at 2% laser power. Envelope analysis was performed using images of sum projections of stacks consisting of 10 frames with custom MATLAB scripts.

To capture kinesin motility dynamics, motility reaction mixtures containing 3 ul of KIF5C-JFX or KIF1A-JFX motor-containing cell extracts were mixed with GFP-tau cell extracts diluted 2-fold in MAB supplemented with 0.2 mg/ml BSA, 10 mM DTT, 20 mM Taxol, 15 mg/ml glucose, ≥ 2000 units/g glucose oxidase, ≥ 6 units/g catalase, 1 mg/ml casein, and 5mM MgATP and flown through the chamber containing microtubules and imaged in TIRF. To image kinesin motility, images were acquired for 2 minutes with 120 msec exposure using 640 nm laser set at 10–15% laser power. Image files were exported as TIFFs, linescans were traced over microtubule images to generate kymographs of kinesin motility, which were then analyzed using KymoButler and custom MATLAB scripts.

### Envelope analysis

To analyze tau envelopes, linescans were traced along microtubules incubated with WT, AP, or E14 GFP-tau, and fluorescence intensity profiles were generated. Tau intensity distributions were then fit using a Gaussian mixture model (GMM) to determine the intensity threshold for defining envelopes. The optimal number of components in each distribution was selected based on the Bayesian Information Criterion (BIC). Microtubules with tau envelopes were identified by multi-component fits, whereas those lacking envelopes were best fit by a single-component model (Fig. 1C). The envelope detection threshold was defined as the intensity corresponding to the first GMM mode plus two standard deviations (σ). Intensities above this threshold were classified as envelopes, while those below were considered diffuse tau between envelopes. A similar analysis was performed to identify and quantify tau envelopes in neurons, with the exception that 10 μm linescans were drawn in envelope-positive regions, which were manually selected based on the degree of fluorescence intensity variation along the axon.

### Analysis of kinesin kinetics on tau-microtubules

To quantify the attachment and detachment frequencies of kinesins on tau-decorated microtubules, tau envelopes were first identified as described above. Kymograph analysis was then used to determine the start and stop positions of each kinesin trajectory, which were categorized as either inside or outside of tau envelopes (Fig. 3G). Pairwise comparisons were performed to assess the significance of differences in attachment and detachment frequencies inside versus outside of envelopes for each tau construct.

To determine the dissociation rate constant of kinesins on tau-decorated microtubules, dwell time distributions were fit to a single exponential function according to equation (2):

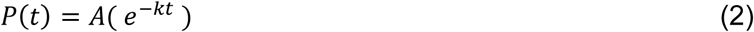

where *P(t)* is the probability of a kinesin detaching at time *t*, *A* is a normalization constant, and *k* is the dissociation rate constant.

### Statistical analysis

All data were presented with error bars indicating either SEM, SD, or 95% CI as specified in the figure legends. All *n* and number of replicates were mentioned in the figure legends. Sample variance significance was determined using a two-sample *F*-test for equal variances in MATLAB. The Wilcoxon signed rank test to test the pairwise comparison of tau intensity in different axonal regions was performed in MATLAB. Bootstrapping analysis was performed in MATLAB and used to test for statistical significance and determine confidence intervals.

## Data availability

Source data files for all figures and extended data figures, are available with this manuscript.

## Acknowledgements

We thank members of the Hendricks lab for helpful discussions and feedback and Dr Gary Brouhard and Dr Kristen J. Verhey for generously providing reagents. This work was supported by NIH.

## Author contributions

D. B., C. L. B., and A. G. H. conceptualization; D. B. formal analysis; D. B. investigation; D. B. and A. G. H. methodology; D. B. and A. G. H. project administration; D. B. and A. G. H. software; D. B. validation; D. B. visualization; D. B. and A. G. H. writing–original draft; D. B., C. L. B., and A. G. H. writing–review and editing; C. L. B. and A. G. H. funding acquisition; C. L. B. and A. G. H. resources; A. G. H. supervision.

## References

Ahmad FJ, Pienkowski TP, Baas PW. Regional differences in microtubule dynamics in the axon. J Neurosci. 1993 Feb;13(2):856–66. doi: 10.1523/JNEUROSCI.13-02-00856.1993. PMID: 8426241; PMCID: PMC6576629.

Alushin GM, Lander GC, Kellogg EH, Zhang R, Baker D, Nogales E. High-resolution microtubule structures reveal the structural transitions in αβ-tubulin upon GTP hydrolysis. Cell. 2014 May 22;157(5):1117–29. doi: 10.1016/j.cell.2014.03.053. PMID: 24855948; PMCID: PMC4054694.

Andreadis A, Brown WM, Kosik KS. Structure and novel exons of the human tau gene. Biochemistry. 1992 Nov 3;31(43):10626–33. doi: 10.1021/bi00158a027. PMID: 1420178.

Arpağ G, Norris SR, Mousavi SI, Soppina V, Verhey KJ, Hancock WO, Tüzel E. Motor Dynamics Underlying Cargo Transport by Pairs of Kinesin-1 and Kinesin-3 Motors. Biophys J. 2019 Mar 19;116(6):1115–1126. doi: 10.1016/j.bpj.2019.01.036. Epub 2019 Feb 5. PMID: 30824116; PMCID: PMC6428962.

Baas PW, Slaughter T, Brown A, Black MM. Microtubule dynamics in axons and dendrites. J Neurosci Res. 1991 Sep;30(1):134–53. doi: 10.1002/jnr.490300115. PMID: 1795398.

Balabanian L, Lessard DV, Swaminathan K, Yaninska P, Sébastien M, Wang S, Stevens PW, Wiseman PW, Berger CL, Hendricks AG. Tau differentially regulates the transport of early endosomes and lysosomes. Mol Biol Cell. 2022 Nov 1;33(13):ar128. doi: 10.1091/mbc.E22-01-0018. Epub 2022 Sep 21. PMID: 36129768; PMCID: PMC9634973.

Beaudet D, Berger CL, Hendricks AG. The types and numbers of kinesins and dyneins transporting endocytic cargoes modulate their motility and response to tau. J Biol Chem. 2024 Jun;300(6):107323. doi: 10.1016/j.jbc.2024.107323. Epub 2024 Apr 25. PMID: 38677516; PMCID: PMC11130734.

Bentley M, Decker H, Luisi J, Banker G. A novel assay reveals preferential binding between Rabs, kinesins, and specific endosomal subpopulations. J Cell Biol. 2015 Feb 2;208(3):273–81. doi: 10.1083/jcb.201408056. Epub 2015 Jan 26. PMID: 25624392; PMCID: PMC4315250.

Binder LI, Frankfurter A, Rebhun LI. The distribution of tau in the mammalian central nervous system. J Cell Biol. 1985 Oct;101(4):1371–8. doi: 10.1083/jcb.101.4.1371. PMID: 3930508; PMCID: PMC2113928.

Biswas S, Kalil K. The Microtubule-Associated Protein Tau Mediates the Organization of Microtubules and Their Dynamic Exploration of Actin-Rich Lamellipodia and Filopodia of Cortical Growth Cones. J Neurosci. 2018 Jan 10;38(2):291–307. doi: 10.1523/JNEUROSCI.2281-17.2017. Epub 2017 Nov 22. PMID: 29167405; PMCID: PMC5761611.

Black MM, Cochran JM, Kurdyla JT. Solubility properties of neuronal tubulin: evidence for labile and stable microtubules. Brain Res. 1984 Mar 19;295(2):255–63. doi: 10.1016/0006-8993(84)90974-0. PMID: 6713187.

Brady ST, Tytell M, Lasek RJ. Axonal tubulin and axonal microtubules: biochemical evidence for cold stability. J Cell Biol. 1984 Nov;99(5):1716–24. doi: 10.1083/jcb.99.5.1716. PMID: 6490717; PMCID: PMC2113352.

Brown CL, Maier KC, Stauber T, Ginkel LM, Wordeman L, Vernos I, Schroer TA. Kinesin-2 is a motor for late endosomes and lysosomes. Traffic. 2005 Dec;6(12):1114–24. doi: 10.1111/j.1600-0854.2005.00347.x. PMID: 16262723.

Budaitis BG, Badieyan S, Yue Y, Blasius TL, Reinemann DN, Lang MJ, Cianfrocco MA, Verhey KJ. A kinesin-1 variant reveals motor-induced microtubule damage in cells. Curr Biol. 2022 Jun 6;32(11):2416–2429.e6. doi: 10.1016/j.cub.2022.04.020. Epub 2022 May 2. PMID: 35504282; PMCID: PMC9993403.

Cario A, Savastano A, Wood NB, Liu Z, Previs MJ, Hendricks AG, Zweckstetter M, Berger CL. The pathogenic R5L mutation disrupts formation of Tau complexes on the microtubule by altering local N-terminal structure. Proc Natl Acad Sci U S A. 2022 Feb 15;119(7):e2114215119.

Chakraborty P, Ibáñez de Opakua A, Purslow JA, et al. GSK3β phosphorylation catalyzes the aggregation of tau into Alzheimer’s disease-like filaments. Proc Natl Acad Sci U S A. 2024;121(52):e2414176121. doi:10.1073/pnas.2414176121

Chaudhary AR, Berger F, Berger CL, Hendricks AG. Tau directs intracellular trafficking by regulating the forces exerted by kinesin and dynein teams. Traffic. 2018 Feb;19(2):111–121. doi: 10.1111/tra.12537. Epub 2017 Dec 5. PMID: 29077261; PMCID: PMC5783771.

Chen J, Kanai Y, Cowan NJ, Hirokawa N. Projection domains of MAP2 and tau determine spacings between microtubules in dendrites and axons. Nature. 1992 Dec 17;360(6405):674–7. doi: 10.1038/360674a0. PMID: 1465130.

Cho JH, Johnson GV. Primed phosphorylation of tau at Thr231 by glycogen synthase kinase 3beta (GSK3beta) plays a critical role in regulating tau’s ability to bind and stabilize microtubules. J Neurochem. 2004 Jan;88(2):349–58. doi: 10.1111/j.1471-4159.2004.02155.x. PMID: 14690523.

Chung PJ, Choi MC, Miller HP, Feinstein HE, Raviv U, Li Y, Wilson L, Feinstein SC, Safinya CR. Direct force measurements reveal that protein Tau confers short-range attractions and isoform-dependent steric stabilization to microtubules. Proc Natl Acad Sci U S A. 2015 Nov 24;112(47):E6416-25. doi: 10.1073/pnas.1513172112. Epub 2015 Nov 5. PMID: 26542680; PMCID: PMC4664379.

Dixit R, Ross JL, Goldman YE, Holzbaur EL. Differential regulation of dynein and kinesin motor proteins by tau. Science. 2008 Feb 22;319(5866):1086–9. doi: 10.1126/science.1152993. Epub 2008 Jan 17. PMID: 18202255; PMCID: PMC2866193.

Fan X, Okada K, Lin H, Ori-McKenney KM, McKenney RJ. A pathological phosphorylation pattern enhances tau cooperativity on microtubules and facilitates tau filament assembly. bioRxiv 2025.01.29.635117; doi: 10.1101/2025.01.29.635117.

Feng Q, Mickolajczyk KJ, Chen GY, Hancock WO. Motor Reattachment Kinetics Play a Dominant Role in Multimotor-Driven Cargo Transport. Biophys J. 2018 Jan 23;114(2):400–409. doi: 10.1016/j.bpj.2017.11.016. PMID: 29401437; PMCID: PMC5985011.

Fulga TA, Elson-Schwab I, Khurana V, Steinhilb ML, Spires TL, Hyman BT, Feany MB. Abnormal bundling and accumulation of F-actin mediates tau-induced neuronal degeneration in vivo. Nat Cell Biol. 2007 Feb;9(2):139–48. doi: 10.1038/ncb1528. Epub 2006 Dec 24. PMID: 17187063.

Goedert M, Wischik CM, Crowther RA, Walker JE, Klug A. Cloning and sequencing of the cDNA encoding a core protein of the paired helical filament of Alzheimer disease: identification as the microtubule-associated protein tau. Proc Natl Acad Sci U S A. 1988 Jun;85(11):4051–5. doi: 10.1073/pnas.85.11.4051. PMID: 3131773; PMCID: PMC280359.

Goedert M, Spillantini MG, Potier MC, Ulrich J, Crowther RA. Cloning and sequencing of the cDNA encoding an isoform of microtubule-associated protein tau containing four tandem repeats: differential expression of tau protein mRNAs in human brain. EMBO J. 1989a Feb;8(2):393–9. doi: 10.1002/j.1460-2075.1989.tb03390.x. PMID: 2498079; PMCID: PMC400819.

Goedert M, Spillantini MG, Jakes R, Rutherford D, Crowther RA. Multiple isoforms of human microtubule-associated protein tau: sequences and localization in neurofibrillary tangles of Alzheimer’s disease. Neuron. 1989b Oct;3(4):519–26. doi: 10.1016/0896-6273(89)90210-9. PMID: 2484340.

Hallinan GI, Vargas-Caballero M, West J, Deinhardt K. Tau Misfolding Efficiently Propagates between Individual Intact Hippocampal Neurons. J Neurosci. 2019;39(48):9623–9632. doi:10.1523/JNEUROSCI.1590-19.2019

Hanger DP, Byers HL, Wray S, et al. Novel phosphorylation sites in tau from Alzheimer brain support a role for casein kinase 1 in disease pathogenesis. J Biol Chem. 2007;282(32):23645–23654. doi:10.1074/jbc.M703269200

Hanger DP, Anderton BH, Noble W. Tau phosphorylation: the therapeutic challenge for neurodegenerative disease. Trends Mol Med. 2009;15(3):112–119. doi:10.1016/j.molmed.2009.01.003

Hasegawa M, Smith MJ, Goedert M. Tau proteins with FTDP-17 mutations have a reduced ability to promote microtubule assembly. FEBS Lett. 1998 Oct 23;437(3):207–10.

Hinrichs MH, Jalal A, Brenner B, Mandelkow E, Kumar S, Scholz T. Tau protein diffuses along the microtubule lattice. J Biol Chem. 2012 Nov 9;287(46):38559–68. doi: 10.1074/jbc.M112.369785. Epub 2012 Sep 27. PMID: 23019339; PMCID: PMC3493901.

Hoeprich GJ, Thompson AR, McVicker DP, Hancock WO, Berger CL. Kinesin’s neck-linker determines its ability to navigate obstacles on the microtubule surface. Biophys J. 2014 Apr 15;106(8):1691–700. doi: 10.1016/j.bpj.2014.02.034. PMID: 24739168; PMCID: PMC4008791.

Hoeprich GJ, Mickolajczyk KJ, Nelson SR, Hancock WO, Berger CL. The axonal transport motor kinesin-2 navigates microtubule obstacles via protofilament switching. Traffic. 2017 May;18(5):304–314. doi: 10.1111/tra.12478. Epub 2017 Apr 5. PMID: 28267259; PMCID: PMC5687255.

Hoover BR, Reed MN, Su J, Penrod RD, Kotilinek LA, Grant MK, Pitstick R, Carlson GA, Lanier LM, Yuan LL, Ashe KH, Liao D. Tau mislocalization to dendritic spines mediates synaptic dysfunction independently of neurodegeneration. Neuron. 2010 Dec 22;68(6):1067–81. doi: 10.1016/j.neuron.2010.11.030. PMID: 21172610; PMCID: PMC3026458.

Iovino M, Patani R, Watts C, Chandran S, Spillantini MG. Human stem cell-derived neurons: a system to study human tau function and dysfunction. PLoS One. 2010 Nov 11;5(11):e13947. doi: 10.1371/journal.pone.0013947. PMID: 21085657; PMCID: PMC2978712.

Iovino M, Agathou S, González-Rueda A, Del Castillo Velasco-Herrera M, Borroni B, Alberici A, Lynch T, O’Dowd S, Geti I, Gaffney D, Vallier L, Paulsen O, Káradóttir RT, Spillantini MG. Early maturation and distinct tau pathology in induced pluripotent stem cell-derived neurons from patients with MAPT mutations. Brain. 2015 Nov;138(Pt 11):3345–59. doi: 10.1093/brain/awv222. Epub 2015 Jul 27. PMID: 26220942; PMCID: PMC4620511.

Jenkins B, Decker H, Bentley M, Luisi J, Banker G. A novel split kinesin assay identifies motor proteins that interact with distinct vesicle populations. J Cell Biol. 2012;198(4):749–761. doi:10.1083/jcb.201205070

Jordens I, Fernandez-Borja M, Marsman M, Dusseljee S, Janssen L, Calafat J, Janssen H, Wubbolts R, Neefjes J. The Rab7 effector protein RILP controls lysosomal transport by inducing the recruitment of dynein-dynactin motors. Curr Biol. 2001 Oct 30;11(21):1680–5. doi: 10.1016/s0960-9822(01)00531-0. PMID: 11696325.

Kanaan NM, Grabinski T. Neuronal and Glial Distribution of Tau Protein in the Adult Rat and Monkey. Front Mol Neurosci. 2021 Apr 27;14:607303. doi: 10.3389/fnmol.2021.607303. PMID: 33986642; PMCID: PMC8112591.

Kanaan NM, Morfini G, Pigino G, LaPointe NE, Andreadis A, Song Y, Leitman E, Binder LI, Brady ST. Phosphorylation in the amino terminus of tau prevents inhibition of anterograde axonal transport. Neurobiol Aging. 2012 Apr;33(4):826.e15-30. doi: 10.1016/j.neurobiolaging.2011.06.006. Epub 2011 Jul 27. PMID: 21794954; PMCID: PMC3272324.

Kanai Y, Chen J, Hirokawa N. Microtubule bundling by tau proteins in vivo: analysis of functional domains. EMBO J. 1992 Nov;11(11):3953–61. doi: 10.1002/j.1460-2075.1992.tb05489.x. PMID: 1396588; PMCID: PMC556906.

Kellogg EH, Hejab NMA, Poepsel S, Downing KH, DiMaio F, Nogales E. Near-atomic model of microtubule-tau interactions. Science. 2018 Jun 15;360(6394):1242–1246. doi: 10.1126/science.aat1780. Epub 2018 May 10. PMID: 29748322; PMCID: PMC6225777.

Li Y, Black MM. Microtubule assembly and turnover in growing axons. J Neurosci. 1996 Jan 15;16(2):531–44. doi: 10.1523/JNEUROSCI.16-02-00531.1996. PMID: 8551337; PMCID: PMC6578637.

Lim SS, Sammak PJ, Borisy GG. Progressive and spatially differentiated stability of microtubules in developing neuronal cells. J Cell Biol. 1989 Jul;109(1):253–63. doi: 10.1083/jcb.109.1.253. PMID: 2745551; PMCID: PMC2115470.

Lindwall G, Cole RD. Phosphorylation affects the ability of tau protein to promote microtubule assembly. J Biol Chem. 1984 Apr 25;259(8):5301–5. PMID: 6425287.

Loubéry S, Wilhelm C, Hurbain I, Neveu S, Louvard D, Coudrier E. Different microtubule motors move early and late endocytic compartments. Traffic. 2008 Apr;9(4):492–509. doi: 10.1111/j.1600-0854.2008.00704.x. Epub 2008 Jan 10. PMID: 18194411.

Mandelkow EM, Biernat J, Drewes G, Gustke N, Trinczek B, Mandelkow E. Tau domains, phosphorylation, and interactions with microtubules. Neurobiol Aging. 1995 May-Jun;16(3):355–62; discussion 362-3. doi: 10.1016/0197-4580(95)00025-a. PMID: 7566345.

Mandelkow EM, Stamer K, Vogel R, Thies E, Mandelkow E. Clogging of axons by tau, inhibition of axonal traffic and starvation of synapses. Neurobiol Aging. 2003 Dec;24(8):1079–85.

McVicker DP, Chrin LR, Berger CL. The nucleotide-binding state of microtubules modulates kinesin processivity and the ability of Tau to inhibit kinesin-mediated transport. J Biol Chem. 2011 Dec 16;286(50):42873–80. doi: 10.1074/jbc.M111.292987. Epub 2011 Oct 27. PMID: 22039058; PMCID: PMC3234877.

McVicker DP, Hoeprich GJ, Thompson AR, Berger CL. Tau interconverts between diffusive and stable populations on the microtubule surface in an isoform and lattice specific manner. Cytoskeleton (Hoboken). 2014 Mar;71(3):184–94. doi: 10.1002/cm.21163. Epub 2014 Feb 24. PMID: 24520046; PMCID: PMC4154625.

Monroy BY, Sawyer DL, Ackermann BE, Borden MM, Tan TC, Ori-McKenney KM. Competition between microtubule-associated proteins directs motor transport. Nat Commun. 2018 Apr 16;9(1):1487. doi: 10.1038/s41467-018-03909-2. PMID: 29662074; PMCID: PMC5902456.

Monroy BY, Tan TC, Oclaman JM, Han JS, Simó S, Niwa S, Nowakowski DW, McKenney RJ, Ori-McKenney KM. A Combinatorial MAP Code Dictates Polarized Microtubule Transport. Dev Cell. 2020 Apr 6;53(1):60–72.e4. doi: 10.1016/j.devcel.2020.01.029. Epub 2020 Feb 27. PMID: 32109385; PMCID: PMC7181406.

Moretto E, Masato A, Panzi C, et al. Aberrant tau accumulation caused by MAPT mutations induces early pathological changes in axonal transport that are rescued by p38α inhibition. Nat Neurosci. 2026;29(6):1355–1368. doi:10.1038/s41593-026-02266-4

Morishima-Kawashima M, Hasegawa M, Takio K, et al. Proline-directed and non-proline-directed phosphorylation of PHF-tau. J Biol Chem. 1995;270(2):823–829. doi:10.1074/jbc.270.2.823

Müller MJ, Klumpp S, Lipowsky R. Tug-of-war as a cooperative mechanism for bidirectional cargo transport by molecular motors. Proc Natl Acad Sci U S A. 2008;105(12):4609–4614. doi:10.1073/pnas.0706825105

Nagpal S, Swaminathan K, Beaudet D, Verdier M, Wang S, Berger CL, Berger F, Hendricks AG. Optogenetic control of kinesin-1, -2, -3 and dynein reveals their specific roles in vesicular transport. Cell Rep. 2024 Aug 27;43(8):114649. doi: 10.1016/j.celrep.2024.114649. Epub 2024 Aug 18. PMID: 39159044; PMCID: PMC11416726.

Niewidok B, Igaev M, Sündermann F, Janning D, Bakota L, Brandt R. Presence of a carboxy-terminal pseudorepeat and disease-like pseudohyperphosphorylation critically influence tau’s interaction with microtubules in axon-like processes. Mol Biol Cell. 2016 Nov 7;27(22):3537–3549. doi: 10.1091/mbc.E16-06-0402. Epub 2016 Aug 31. PMID: 27582388; PMCID: PMC5221586.

Okabe S, Hirokawa N. Turnover of fluorescently labelled tubulin and actin in the axon. Nature. 1990 Feb 1;343(6257):479–82. doi: 10.1038/343479a0. PMID: 1689016.

Panda D, Samuel JC, Massie M, Feinstein SC, Wilson L. Differential regulation of microtubule dynamics by three- and four-repeat tau: implications for the onset of neurodegenerative disease. Proc Natl Acad Sci U S A. 2003 Aug 5;100(16):9548–53. doi: 10.1073/pnas.1633508100. Epub 2003 Jul 28. PMID: 12886013; PMCID: PMC170955.

Piras A, Collin L, Grüninger F, Graff C, Rönnbäck A. Autophagic and lysosomal defects in human tauopathies: analysis of post-mortem brain from patients with familial Alzheimer disease, corticobasal degeneration and progressive supranuclear palsy. Acta Neuropathol Commun. 2016 Mar 2;4:22.

Pu J, Guardia CM, Keren-Kaplan T, Bonifacino JS. Mechanisms and functions of lysosome positioning. J Cell Sci. 2016 Dec 1;129(23):4329–4339. doi: 10.1242/jcs.196287. Epub 2016 Oct 31. PMID: 27799357; PMCID: PMC5201012.

Rodríguez-Martín T, Pooler AM, Lau DHW, Mórotz GM, De Vos KJ, Gilley J, Coleman MP, Hanger DP. Reduced number of axonal mitochondria and tau hypophosphorylation in mouse P301L tau knockin neurons. Neurobiol Dis. 2016 Jan;85:1–10.

Rosa-Ferreira C, Munro S. Arl8 and SKIP act together to link lysosomes to kinesin-1. Dev Cell. 2011 Dec 13;21(6):1171–8. doi: 10.1016/j.devcel.2011.10.007. PMID: 22172677; PMCID: PMC3240744.

Rosenberg KJ, Ross JL, Feinstein HE, Feinstein SC, Israelachvili J. Complementary dimerization of microtubule-associated tau protein: Implications for microtubule bundling and tau-mediated pathogenesis. Proc Natl Acad Sci U S A. 2008 May 27;105(21):7445–50. doi: 10.1073/pnas.0802036105. Epub 2008 May 21. PMID: 18495933; PMCID: PMC2396711.

Sahenk Z, Brady ST. Axonal tubulin and microtubules: morphologic evidence for stable regions on axonal microtubules. Cell Motil Cytoskeleton. 1987;8(2):155–64. doi: 10.1002/cm.970080207. PMID: 2891447.

Schneider A, Biernat J, von Bergen M, Mandelkow E, Mandelkow EM. Phosphorylation that detaches tau protein from microtubules (Ser262, Ser214) also protects it against aggregation into Alzheimer paired helical filaments. Biochemistry. 1999 Mar 23;38(12):3549–58. doi: 10.1021/bi981874p. PMID: 10090741.

Siahaan V, Krattenmacher J, Hyman AA, Diez S, Hernández-Vega A, Lansky Z, Braun M. Kinetically distinct phases of tau on microtubules regulate kinesin motors and severing enzymes. Nat Cell Biol. 2019 Sep;21(9):1086–1092. doi: 10.1038/s41556-019-0374-6. Epub 2019 Sep 2. PMID: 31481789.

Siahaan V, Tan R, Humhalova T, Libusova L, Lacey SE, Tan T, Dacy M, Ori-McKenney KM, McKenney RJ, Braun M, Lansky Z. Microtubule lattice spacing governs cohesive envelope formation of tau family proteins. Nat Chem Biol. 2022 Nov;18(11):1224–1235. doi: 10.1038/s41589-022-01096-2. Epub 2022 Aug 22. PMID: 35996000; PMCID: PMC9613621.

Siahaan V, Weissova R, Karhanova A, et al. Tau phosphorylation impedes functionality of protective tau envelopes. Nat Chem Biol. 2026;22(5):759–769. doi:10.1038/s41589-025-02122-9

Song Y, Kirkpatrick LL, Schilling AB, Helseth DL, Chabot N, Keillor JW, Johnson GV, Brady ST. Transglutaminase and polyamination of tubulin: posttranslational modification for stabilizing axonal microtubules. Neuron. 2013 Apr 10;78(1):109–23. doi: 10.1016/j.neuron.2013.01.036. PMID: 23583110; PMCID: PMC3627183.

Sposito T, Preza E, Mahoney CJ, Setó-Salvia N, Ryan NS, Morris HR, Arber C, Devine MJ, Houlden H, Warner TT, Bushell TJ, Zagnoni M, Kunath T, Livesey FJ, Fox NC, Rossor MN, Hardy J, Wray S. Developmental regulation of tau splicing is disrupted in stem cell-derived neurons from frontotemporal dementia patients with the 10 + 16 splice-site mutation in MAPT. Hum Mol Genet. 2015 Sep 15;24(18):5260–9. doi: 10.1093/hmg/ddv246. Epub 2015 Jul 1. PMID: 26136155; PMCID: PMC4550814.

Stamer K, Vogel R, Thies E, Mandelkow E, Mandelkow EM. Tau blocks traffic of organelles, neurofilaments, and APP vesicles in neurons and enhances oxidative stress. J Cell Biol. 2002 Mar 18;156(6):1051–63. doi: 10.1083/jcb.200108057. Epub 2002 Mar 18. PMID: 11901170; PMCID: PMC2173473.

Steinhilb ML, Dias-Santagata D, Mulkearns EE, Shulman JM, Biernat J, Mandelkow EM, Feany MB. S/P and T/P phosphorylation is critical for tau neurotoxicity in Drosophila. J Neurosci Res. 2007 May 1;85(6):1271–8. doi: 10.1002/jnr.21232. PMID: 17335084.

Stern JL, Lessard DV, Hoeprich GJ, Morfini GA, Berger CL. Phosphoregulation of Tau modulates inhibition of kinesin-1 motility. Mol Biol Cell. 2017 Apr 15;28(8):1079–1087. doi: 10.1091/mbc.E16-10-0728. Epub 2017 Mar 1. PMID: 28251926; PMCID: PMC5391184.

Stokin GB, Lillo C, Falzone TL, Brusch RG, Rockenstein E, Mount SL, Raman R, Davies P, Masliah E, Williams DS, Goldstein LS. Axonopathy and transport deficits early in the pathogenesis of Alzheimer’s disease. Science. 2005 Feb 25;307(5713):1282–8.

Tan R, Lam AJ, Tan T, Han J, Nowakowski DW, Vershinin M, Simó S, Ori-McKenney KM, McKenney RJ. Microtubules gate tau condensation to spatially regulate microtubule functions. Nat Cell Biol. 2019 Sep;21(9):1078–1085. doi: 10.1038/s41556-019-0375-5. Epub 2019 Sep 2. PMID: 31481790; PMCID: PMC6748660.

Trinczek B, Biernat J, Baumann K, Mandelkow EM, Mandelkow E. Domains of tau protein, differential phosphorylation, and dynamic instability of microtubules. Mol Biol Cell. 1995 Dec;6(12):1887–902. doi: 10.1091/mbc.6.12.1887. PMID: 8590813; PMCID: PMC366657.

Vershinin M, Carter BC, Razafsky DS, King SJ, Gross SP. Multiple-motor based transport and its regulation by Tau. Proc Natl Acad Sci U S A. 2007 Jan 2;104(1):87–92. doi: 10.1073/pnas.0607919104. Epub 2006 Dec 26. PMID: 17190808; PMCID: PMC1765483.

Vershinin M, Xu J, Razafsky DS, King SJ, Gross SP. Tuning microtubule-based transport through filamentous MAPs: the problem of dynein. Traffic. 2008 Jun;9(6):882–92. doi: 10.1111/j.1600-0854.2008.00741.x. Epub 2008 Mar 28. PMID: 18373727; PMCID: PMC2958055.

Weingarten MD, Lockwood AH, Hwo SY, Kirschner MW. A protein factor essential for microtubule assembly. Proc Natl Acad Sci U S A. 1975 May;72(5):1858–62. doi: 10.1073/pnas.72.5.1858. PMID: 1057175; PMCID: PMC432646.

Wesseling H, Mair W, Kumar M, et al. Tau PTM Profiles Identify Patient Heterogeneity and Stages of Alzheimer’s Disease. Cell. 2020;183(6):1699–1713.e13. doi:10.1016/j.cell.2020.10.029

