## Supplemental Figures for "Tau hyperphosphorylation impairs cooperative binding to microtubules and perturbs organelle trafficking in neurons"

**Supporting Information**

This PDF file includes:

Supplemental Figures S1 to S4

Supplemental Table S1 and S2

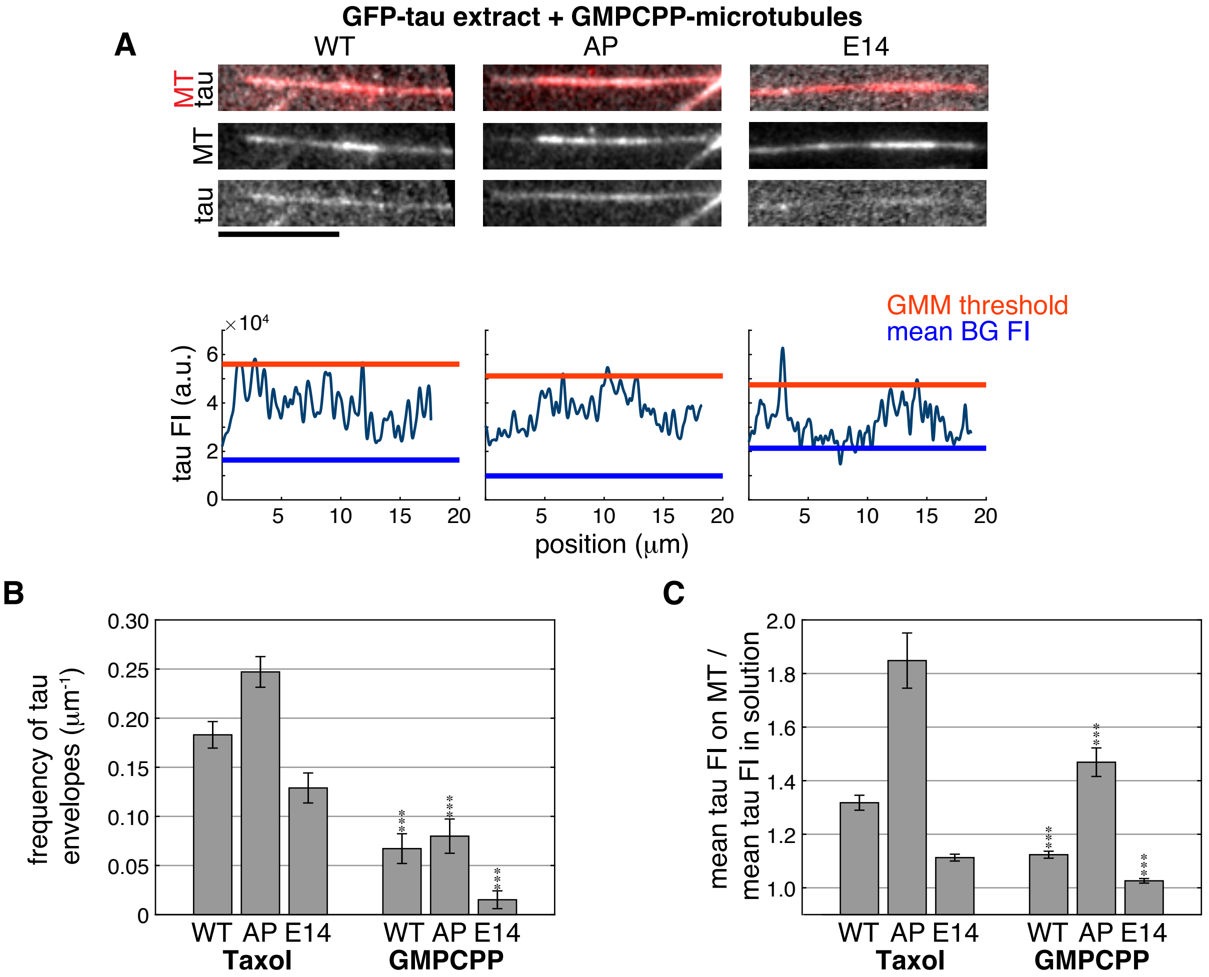

GFP-tau extract + GMPCPP-microtubules
A
WT
AP
E14
MT
tau
MT
tau
GMM threshold
mean BG FI
×10⁴
tau FI (a.u.)
position (μm)
B
frequency of tau envelopes (μm⁻¹)
WT AP E14
Taxol
WT AP E14
GMPCPP
C
mean tau FI on MT / mean tau FI in solution
WT AP E14
Taxol
WT AP E14
GMPCPP


**Figure S1**

**Figure S1.** Tau hyperphosphorylation results in more diffuse microtubule interactions
**A)** Representative images of WT, AP, and E14 GFP-tau on GMPCPP-polymerized microtubules (WT: *n*=180, AP: *n*=178, E14: *n*=184). The indicated *n* values represent total number of samples over 3–4 replicates. Below the images, corresponding tau intensity plots along microtubules are shown, with horizontal lines indicating the GMM threshold (orange) and mean background fluorescence intensity (blue). **B–C)** Bar plots compare the effects of tau phosphorylation on envelope frequency (B) and the mean tau intensity on microtubules versus in solution using either reversibly stabilized (taxol) or irreversibly stabilized (GMPCPP) microtubules. Error bars indicate 95% CI. Statistical significance was assessed using Student's t-test (*** $p < 0.0001$). Scale bars are 10 μm.

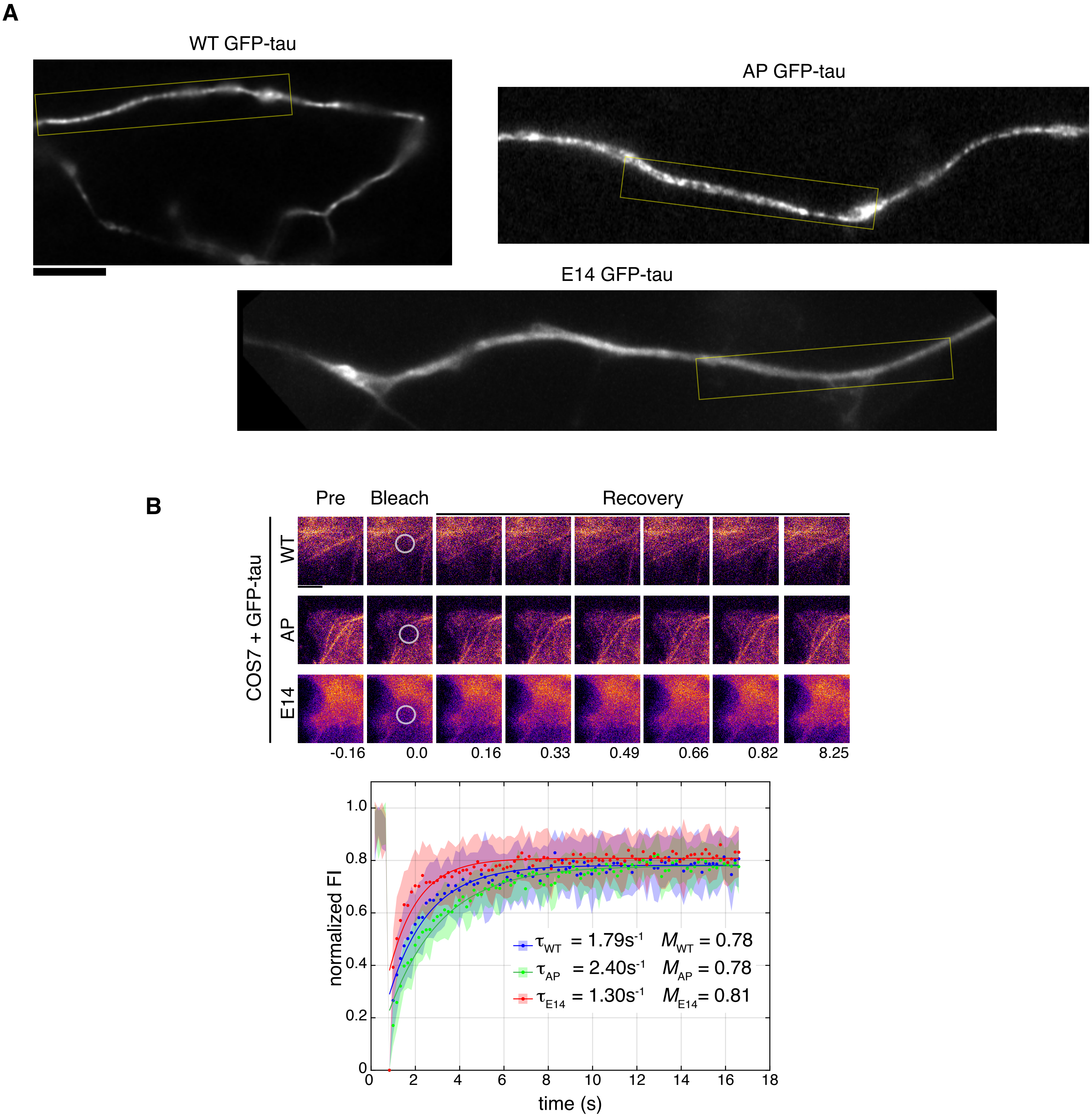
A
WT GFP-tau
AP GFP-tau
E14 GFP-tau
B
Pre
Bleach
Recovery
COS7 + GFP-tau
WT
AP
E14
-0.16
0.0
0.16
0.33
0.49
0.66
0.82
8.25
normalized FI
time (s)
$\tau_{WT}$ = 1.79s$^{-1}$ $M_{WT}$ = 0.78
$\tau_{AP}$ = 2.40s$^{-1}$ $M_{AP}$ = 0.78
$\tau_{E14}$ = 1.30s$^{-1}$ $M_{E14}$= 0.81

**Figure S2**

**Figure S2.** Characterization of GFP-tau phospho-variants in neurons and COS-7 cells
**A)** Full-field images of MAPT-KO neurons expressing WT, AP, and E14 GFP-tau. Yellow boxes indicate the axonal regions shown as cropped images in Fig. 2A. **B)** Timelapse images of fluorescence recovery after photobleaching assays of COS-7 cells expressing WT($n$=15), AP($n$=16), or E14 ($n$=15) GFP-tau. Circles indicate the bleached regions. Fluorescence recovery curves of WT (blue), AP (green), and E14 tau (red). Shaded regions indicate SD. The characteristic recovery ($\tau$) and mobile fraction ($M$) are indicated on the plot for each tau construct. Scale bars are 10 μm (A) and 5 μm (B).

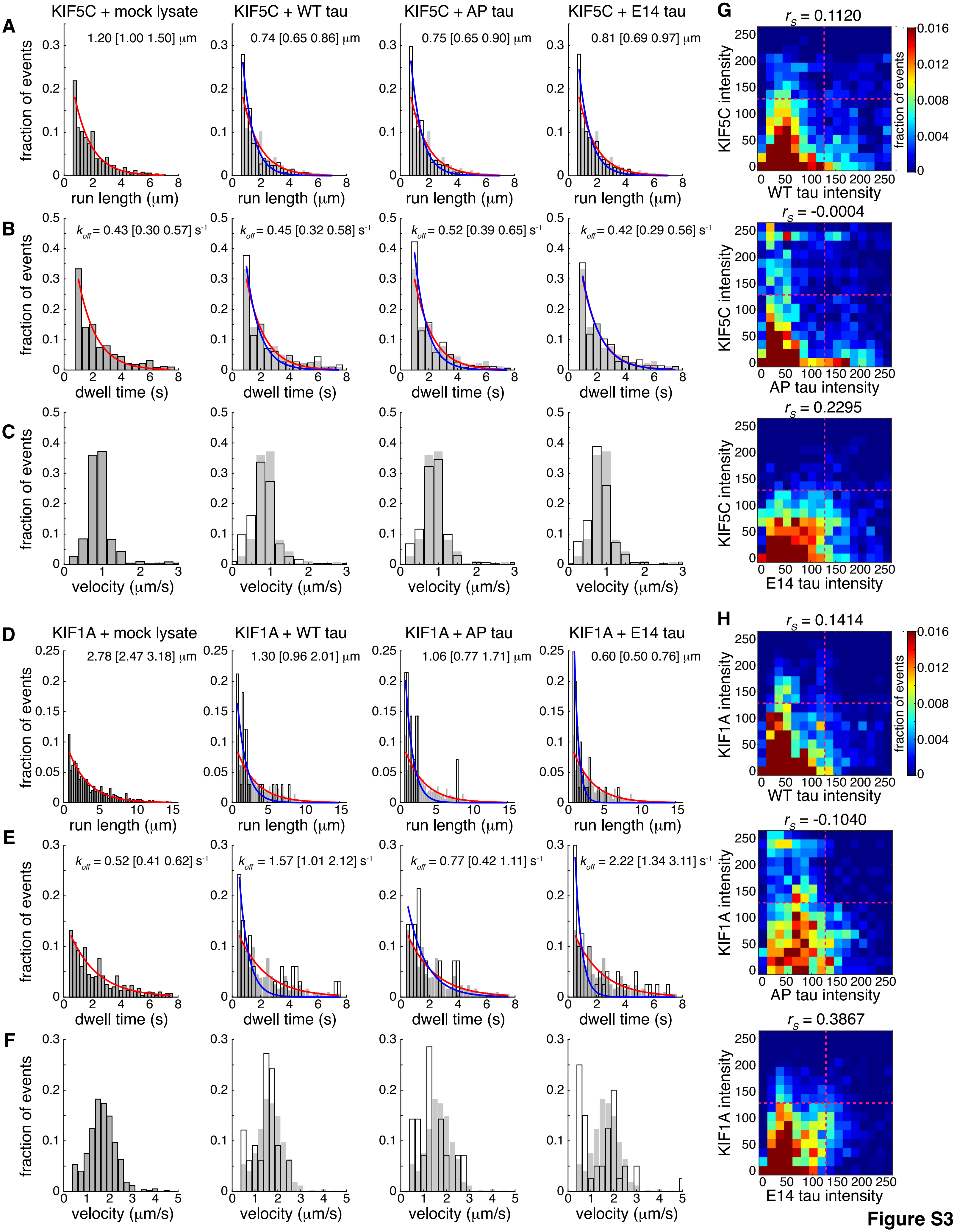

KIF5C + mock lysate
KIF5C + WT tau
KIF5C + AP tau
KIF5C + E14 tau
A
1.20 [1.00 1.50] μm
0.74 [0.65 0.86] μm
0.75 [0.65 0.90] μm
0.81 [0.69 0.97] μm
fraction of events
run length (μm)
B
$k_{off}$ = 0.43 [0.30 0.57] $s^{-1}$
$k_{off}$ = 0.45 [0.32 0.58] $s^{-1}$
$k_{off}$ = 0.52 [0.39 0.65] $s^{-1}$
$k_{off}$ = 0.42 [0.29 0.56] $s^{-1}$
dwell time (s)
C
velocity (μm/s)
G
$r_s$ = 0.1120
KIF5C intensity
WT tau intensity
fraction of events
$r_s$ = -0.0004
AP tau intensity
$r_s$ = 0.2295
E14 tau intensity
KIF1A + mock lysate
KIF1A + WT tau
KIF1A + AP tau
KIF1A + E14 tau
D
2.78 [2.47 3.18] μm
1.30 [0.96 2.01] μm
1.06 [0.77 1.71] μm
0.60 [0.50 0.76] μm
E
$k_{off}$ = 0.52 [0.41 0.62] $s^{-1}$
$k_{off}$ = 1.57 [1.01 2.12] $s^{-1}$
$k_{off}$ = 0.77 [0.42 1.11] $s^{-1}$
$k_{off}$ = 2.22 [1.34 3.11] $s^{-1}$
F
H
$r_s$ = 0.1414
KIF1A intensity
$r_s$ = -0.1040
$r_s$ = 0.3867


**Figure S3**

**Figure S3.** Tau phosphorylation differentially regulates kinesin-1 and kinesin-3 motility
**A–F)** Histograms display the distributions of run lengths, dwell times, and velocities for KIF5C (A–C) and KIF1A (D–F) in the presence of mock lysate or lysates containing WT, AP, or E14 GFP-tau. Run length distributions (A, D) and dwell time distributions (B, E) were fitted with a single exponential function to extract the characteristic run length and the dissociation rate constants ($k_{off}$), respectively. For visual comparison, distributions from tau-containing conditions are shown as black-outlined transparent bars, overlaid on light grey bars representing the control (mock lysate) distribution. Red lines indicate exponential fits for control conditions, and blue lines represent fits for each tau condition. The extracted run lengths and $k_{off}$ values are displayed next to each plot, with 95% CI shown in brackets. **G and H)** Heat maps show the correlation between tau pixel intensities along microtubules with standard deviation projections of kinesin intensity, where bright pixels indicate higher dynamics. Spearman's correlation analysis was performed showing that there is no relationship between kinesin dynamics and tau intensities ($r_S$ values indicated above each plot). Colors represent the fraction of pixels for each bin. Dashed magenta lines are present to highlight the subtle differences of KIF5C and KIF1A dynamics in regions of higher tau intensities across tau phospho-variants.

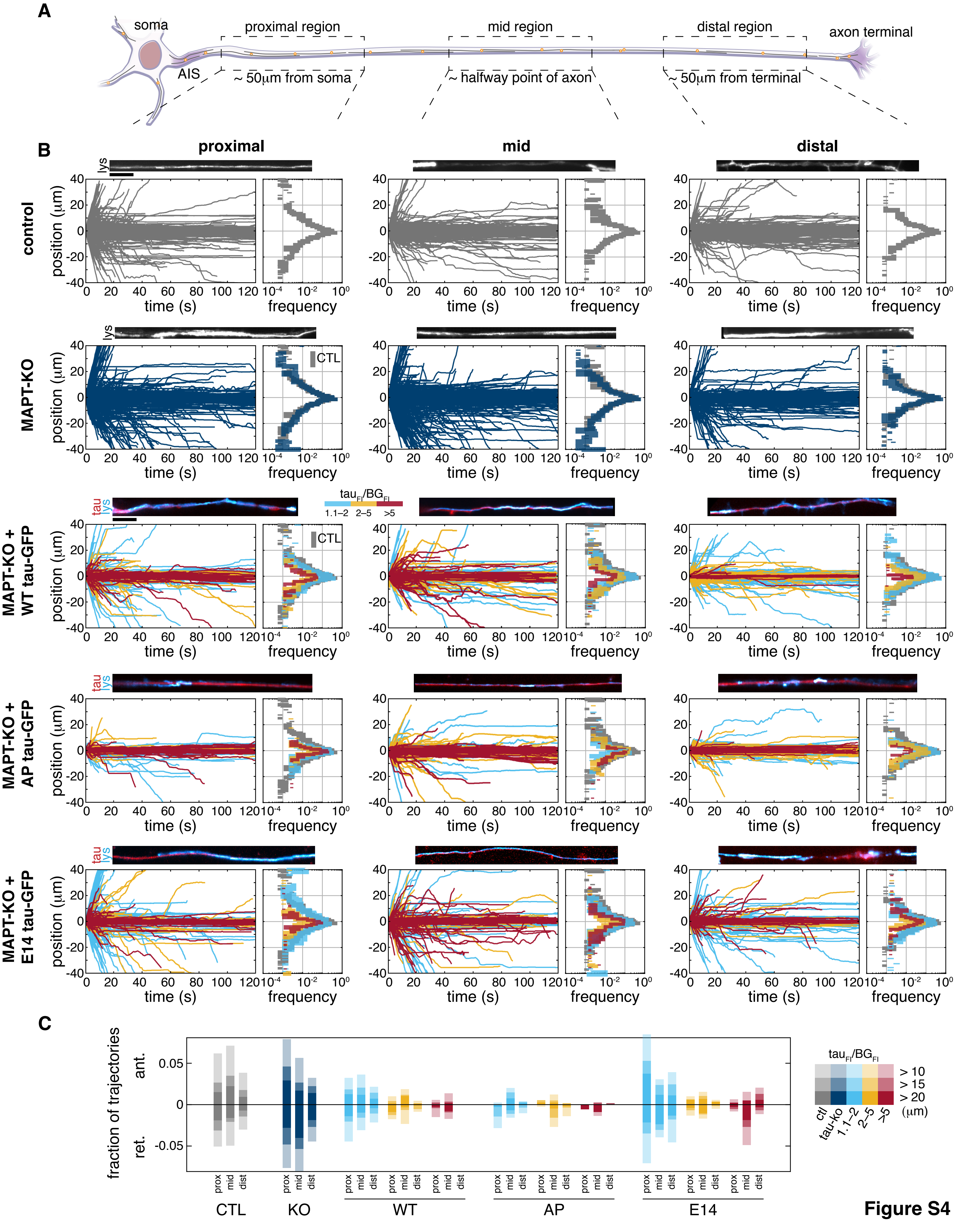
A
soma
AIS
proximal region
~ 50μm from soma
mid region
~ halfway point of axon
distal region
~ 50μm from terminal
axon terminal
B
proximal
mid
distal
control
MAPT-KO
MAPT-KO + WT tau-GFP
MAPT-KO + AP tau-GFP
MAPT-KO + E14 tau-GFP
lys
tau
CTL
tau$_{FI}$/BG$_{FI}$
1.1–2 2–5 >5
position (μm)
time (s)
frequency
C
fraction of trajectories
ant.
ret.
prox mid dist
CTL
KO
WT
AP
E14
ctl tau-ko 1.1–2 2–5 >5
> 10
> 15
> 20
(μm)

Figure S4

**Figure S4A.** Tau perturbations impact the trafficking of lysosomes in different axonal regions
**A)** Schematic of a neuron indicating the regions where tau's effects on lysosome transport were assessed. Lysosome transport was recorded in proximal, mid, and distal axonal regions. Proximal regions were defined as ~50 μm from the soma, mid-axonal regions as ~halfway along the axon, and distal regions as ~50 μm from the axon terminal. Images are all oriented so that the soma is towards the left and the distal axon is towards the right. **B)** Maximum intensity projections show lysosomes (LysoTracker) in the proximal, mid, and distal regions of the axon in DIV 7–8 iPSC-derived control neurons, MAPT-KO neurons, and MAPT-KO neurons expressing WT, AP, or E14 GFP-tau. Images are all oriented so that the soma is towards the left and the distal axon is towards the right. Below each set of images, plots show the total number of lysosome trajectories mapped as position over time. Histograms next to each plot show the frequency distribution of lysosome travel distances for each condition. Positive values indicate anterograde transport (towards distal axon), while negative values indicate retrograde transport (towards soma). Trajectory plots are color-coded based on GFP-tau expression level: light blue (1.1–2× background), orange (2–5×), and red (>5×). Corresponding travel distance histograms are color-coded and overlaid with control data (grey) for comparison. Scale bars are 10 μm. **C)** Bar plot shows how tau expression levels affect the fraction of anterograde and retrograde long-distance trajectories for control (CTL), MAPT-KO (KO), and MAPT-KO neurons expressing WT, AP, or E14 GFP-tau in the proximal, mid, and distal axon.

**Table 1. GFP-tau concentration in cell lysate**

| | ***Concentration (μM) (± SD)*** | ***Percentage of total protein*** |
|---|---|---|
| ***WT GFP-tau*** | **1.10 ± 0.01** | **3.1%** |
| ***AP GFP-tau*** | **1.08 ± 0.13** | **3.3%** |
| ***E14 GFP-tau*** | **1.06 ± 0.04** | **3.1%** |

**Table 2. Overview of *in vivo* lysosome transport data and sample size**

| | ***Proximal*** | ***Mid*** | ***Distal*** |
|---|---|---|---|
| ***Control*** | 3525 trajectories from 38 cells | 2176 trajectories from 30 cells | 3028 trajectories from 44 cells |
| ***MAPT-KO*** | 4709 trajectories from 50 cells | 6416 trajectories from 64 cells | 3571 trajectories from 43 cells |
| ***MAPT-KO + WT tau*** | 1971 trajectories from 31 cells | 1622 trajectories from 26 cells | 2429 trajectories from 41 cells |
| ***MAPT-KO + AP tau*** | 1027 trajectories from 22 cells | 1114 trajectories from 23 cells | 1585 trajectories from 29 cells |
| ***MAPT-KO + E14 tau*** | 2015 trajectories from 23 cells | 1505 trajectories from 23 cells | 1506 trajectories from 26 cells |
